# Functional genome and microbiome in blood of goats affected by the gastrointestinal pathogen *Haemonchus contortus*

**DOI:** 10.1101/2021.04.09.439205

**Authors:** Yonathan Tilahun

## Abstract

The Alpine goat *Capra aegagrus hircus* is parasitized by the barber pole worm (*Haemonchus contortus*). This relationship results in changes that affect the gene expression of the host, the pest, and the microbiome of both. Hematological parameters indicating genes that are expressed and/or the % Composition of abundant and diverse microbial flora are reflective of infestation. We explored the similarity/dissimilarity between and among blood samples of non-infected, infected, infected zoledronic acid-treated, and infected antibody (anti-γσ T cells) treated wethers under controlled conditions. We identified responses to barber pole worms using blood-based analysis of transcripts and the microbiome. Seven (7) days post-inoculation (dpi) we identified 7,627 genes associated with different treatment types. Across all treatments we identified fewer raw read counts and a reduced diversity in microbial flora on 7 dpi than in 21 dpi wethers. We also identified that there were differences in % Composition of microbial flora known to be associated with inflammation. This study identifies treatment specific genes, and an increase in microflora abundance and diversity as wethers age post infestation. Further, *Firmicutes*/*Bacteriodetes* (F/B) ratio reflect metabolic health, based on depression or elevation above thresholds defined by the baseline of non-infected hosts depending on the type of intrusion exhibited by the pest.

## Introduction

*Capra aegagrus hircus* (the Alpine goat) is parasitized by many strongylid nematodes. Among these the barber pole worm (*Haemonchus contortus*) is particularly important [1]. They are the principal parasites of goats, causing global losses to agriculture estimated at over $100 billion each year, increasing every year since 1987 [2]. This nematode is a blood-sucking parasite and can remove < 30 μL of blood/day from the host, inducing anemia and often death.

*Haemonchus contortus* can prove fatal if untreated in cattle, sheep, and goats [3]. Adult worms are inhibited by goats until the environment becomes favorable. Such circumstances increase susceptibility in kids up to eight weeks after parturition [4]. Even moderate infections can reduce milk production and lead to stunted growth in kids. Other effects include anemia, low packed cell volume (PCV), diarrhea, dehydration, peripheral, and internal fluid accumulation, lower reproductive performance, higher mortality and more frequent illness [5]. Worm egg counts are the primary method to diagnose worm infections before localized production losses [6]. Significant blood loss leads to visual signs of infestation that may be confused with or due to a combined effect from other types of parasites and diseases.

A. *H. contortus* produces excretory and secretory products that depress the host immune response [7,8,9]. The host counters through immune responses by its genome and microbiome in a back-and-forth arms race [9]. The specific method that is used to avoid host surveillance is not fully understood, but one theory suggests helminth inflammation inhibition is completed by modulating butyrate biosynthesis [10, 8].

The microbiome is composed of bacteria, archeal, viral, and fungal microbial taxa. These may be commensal, mutualistic, or pathogenic, serving roles ranging from beneficial to inconsequential or detrimental [11]. We described our performance and analysis to identify differences in microbiome composition displaying significant differences in abundance between uninfected control wethers, infected only wethers with *H. contortus*, infected wethers treated with zoledronic acid (ZA), and infected wethers treated with anti-γσ T cell antibodies (AB). These different states or treatments will estimate “metabolic health” in small ruminant goat blood microbiomes (GBMs) following infection by the barber pole nematode *H. contortus*. A large array of products exuded from intestinal helminths modulate microbial community growth and metabolism [11]. Also, *H. contortus* compete with naturally occurring flora of the host for energy-rich nutrients or essential minerals [12]. Infection by *H. contortus* is known to impact intestinal physiology by increasing fluid secretion that alters the habitat of healthy bacterial communities [11]. We examine if this impacts the GBM.

Helminth infestations are known to quickly change the metabolic activity of the abomasum in hosts versus unafflicted small ruminants [7,13,8]. An array of structures, cells, and secretions respond to infestation [14–16]. Natural microbiotas provide resources for innate immunity [16]. This typically happens via altered microbiota diversity richness when compared to un-infested animals [10].

The genes expressed by an organism’s genome are major players in how and when the host responds to a parasite. Additionally, the microbiome modulates responses via internal and external environmental cues [17]. Infestation of a host by a parasite interact with surrounding environmental factors to form an intricate web of stimuli and responses by both entities. The microbiome influences many aspects of these diverse ecosystems [18]. A tri-directional interaction is predicted [19], whereby the host depends on its own genome and its vast microbiome content to defend itself against external stressors including parasite intrusions.

Although blood is assumed to be sterile, devoid of other types of cells [20, 21], microorganisms often occur in blood without inducing disease [20,21,22]. Here, we describe the blood expressed transcriptomes, the abundance and diversity of resident microorganisms, their phylogenetic affiliations, and their relevance towards a metabolically healthy GBM of both uninfected and infected small ruminant *Capra hircus* wethers with *H. contortus*.

We hypothesize that the microflora composition responds to parasite infestation; therefore, the immune response is expressed through genes exhibited by the host genome and its microbiome. We hypothesized that the host genome and the microbiome respond to invasion by parasites by depressing the *Firmicutes*/*Bacteriodetes* ratio blow threshold. We predicted that host genomes would express more immunological genes and that microfloral composition and populations would change over time.

### Objectives

The objectives of the research described here are to identify if blood may be used as a parasite diagnostic tool by: 1) analyzing the different responses to different treatments (uninfected wethers, infected wethers with *H. contortus*, infected wethers treated with zoledronic acid (ZA), and infected wethers treated with anti-γσ T cell antibodies (AB)) as determined by transcriptomic and metagenomic analyses; 2) identifying genes expressed from these treatments after seven (7) days post inoculation (dpi); 3) comparing microbial flora abundance and diversity between seven 7 dpi and 21 dpi using operational taxonomic units (OTUs); 4) and by determining if there are significant differences between *Firmicutes*/*Bacteriodetes* (*F*/*B*) ratios as an indication of infection in order to determine differences of “metabolic health” in wethers.

## Materials and Methods

### Ethical statement

The treatment of animals in our research abided by the guidelines of the Langston University Institutional Animal Care and Use Committee (LUACUC) Approval # 2018-14.

### Animals and treatments

Forty (40) Alpine wethers (114.2±0.92 d of age; 19.4±0.33 kg BW at the start of experiment) being raised in indoor pens at the Langston University farm were used. The animals were checked for fecal egg counts (FECs) and confirmed that they were nematode-parasite free. The animals were allocated randomly to four (4) groups of 10 animals each, and two (2) or three (3) animals from each group were assigned to one of the four (4) pens. All animals were allowed to acclimatize to pens and feeders for daily supplies of 200 g concentrate pellet per animal composed of 500 g of ground grass (50%) and alfalfa (50%) hays. The treatment groups were as:

**Table 1.**
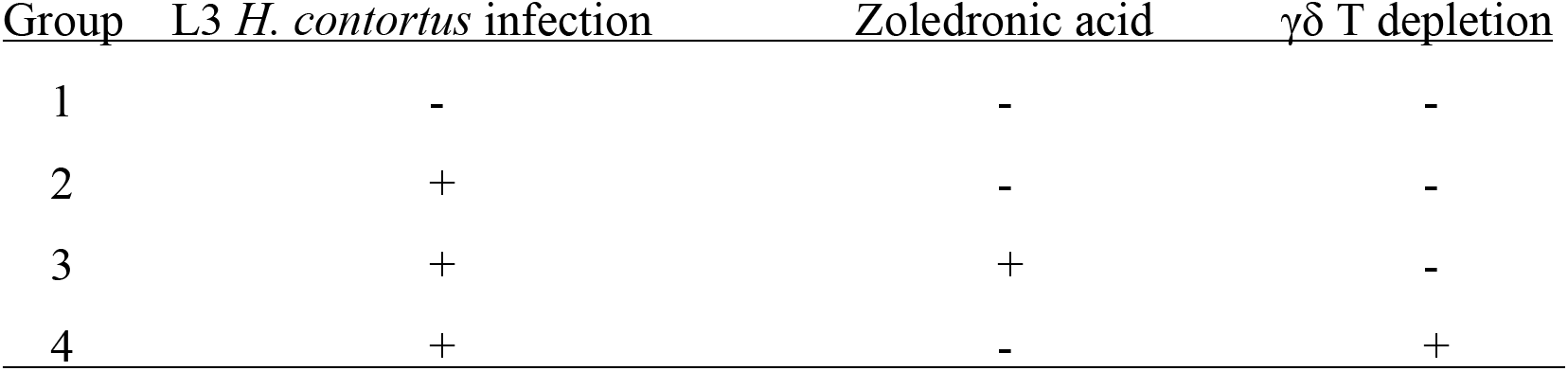
Experimental Set-up. Treatment Group (1-4) for infection L3 *H. contortus* (+) or non-infection *H. contortus* (−), with (+) or without (−) treatment type (Zoledronic acid injection or γδ T depletion).

On the first (1) day prior to the L3 infection, the anti-γδ T cells antibody were administered intravenously. ZA were administered intravenously 7 days prior to and 0, 7, and 14 days after the L3 infection. At the beginning of the experiment all kids except Group 1 were given 10,000 *H. contortus* infective larvae (L3; hatched and isolated from feces being collected from LU goats) by gavage. Five animals from each group were euthanized on seven (7) days post inoculation (dpi) for sampling and the other five (5) animals were euthanized 21 dpi.

### Blood sample collection and processing

Blood samples were collected from five goats in four treatment groups (No infection, Infection, Infection ZA inject, and Infection AB inject), following 7 dpi. Blood samples included red blood cells, white blood cells (total and differential), hemoglobin, platelets, and plasma protein, from the jugular vein in EDTA tubes. Quality assurance/quality control (QA/QC) parameters resulted in blood samples from 19/20 cDNA libraries that were used from samples collected 7 dpi. The cDNA libraries were sequenced on an illumina RNA-Seq Next-generation sequencing (NGS) instrument and filtered and normalized using Partek^®^ Flow^®^ software suites.

Also, methods of identifying naturally occurring microbial flora in nontreated and treated wethers were identified. Analytics for QA/QC for high-throughput barcoded illumina MiSeq NGS sequencing of 16S rRNA, resulted in 18/20 samples that were obtained for 7 dpi and 20/20 samples for 21 dpi.

### Total RNA Purification

Total RNA was collected from blood samples using a modified TRIzol reagent procedure. The blood samples were lysed by using ice chilled TRIzol Reagent. The broken cells were homogenized by pipetting up and down many times. After transferring the homogenized and broken cells into Eppendorf, Chloroform was added to the lysed cells. It gave three layers. The upper aqueous layer contained extracted RNA, an interphase contained DNA, and proteins were dissolved in the bottom red organic layer. The pH was kept at around 4 for RNA purification, which held RNA in the aqueous phase preferentially. After centrifugation, the upper aqueous layer was pipetted out carefully and isopropanol was added to precipitate the RNA. Again, the RNA was precipitated by centrifugation to get RNA pellets which were washed with 70% ethanol (made with DEPC treated water), air-dried and suspended in DEPC treated (RNase free) water. Extracted RNA was quantified to calculate yield by a NanoDrop^TM^ 2000 spectrophotometer.

### A. Lysate Preparation from Blood

The cDNA libraries were constructed by initial first strand synthesis using the Protocol for Non-directional RNA-seq Workflows and NEBNEXT^®^ Ultra^TM^ II RNA First Strand Synthesis Module (E7771), according to a modified manufacturer’s protocol as follows:

### Input Amount Requirement

A 100 ng total RNA was quantified after the purification. The protocol was optimized for approximately 200 nt RNA inserts. The protocol was optimized using Universal Human Reference Total RNA.

### RNA Fragmentation and Priming

The fragmentation and priming reactions were assembled on ice in a nuclease-free tube by adding the Fragmentation and Priming Mix for a total volume of 10 μL. They were mixed thoroughly by pipetting. The samples were placed in a thermocycler and incubated at 94 °C. The tube was immediately transferred to ice and First Strand cDNA Synthesis was begun immediately.

### First Strand cDNA Synthesis Reaction

The first strand synthesis reaction was assembled on ice by adding components to the fragmented and primed RNA for a total volume of 20 μL. The reaction was mixed thoroughly by pipetting. The sample was incubated in a preheated thermocycler with the heated lid set at ≥ 80 ° C as follows: Step 1: 10 minutes at 25° C; Step 2: 15 minutes at 42 °C; Step 3: 15 minutes at 70 ° C; and Step 4: Hold at 4° C. We then proceeded directly to Swift Biosciences^TM^ ACCEL-NGS^®^ 1S PLUS DNA LIBRARY KIT: Single, Dual Combinatorial and Unique Dual Indexing and prepared the DNA Libraries as follows:

### Denaturation

Due to the short incubation time of the denaturation step, all of the reagents of the Adaptase Reaction Mix were pre-assembled and placed on ice. The thermocycler was pre-heated to 95° C. The fragmented DNA sample was transferred to a 0.2 mL PCR tube and the volume of the sample adjusted to a final volume of 15 μL using Low EDTA TE, if it was necessary. The samples were placed in the thermocycler, programmed at 95 ° C for 2 minutes with lid heating ON. Upon completion, the tube(s) were placed on ice immediately for 2 minutes. We then proceeded directly to the Adaptase step to preserve the maximum amount of ssDNA substrate.

### Adaptase

The Adaptase Thermocycler Program was loaded on the thermocycler and paused at the first step to pre-heat to 37 ° C until all samples were loaded. Twenty-five (25) μL of the pre-assembled Adaptase Reaction Mix was added to each PCR tube containing a 15 μL DNA sample and mixed by pipetting or gentle vortexing until homogeneous, after which they were spun down. The samples in the thermocycler were run at the following parameters with the lid heating ON. The thermocycler Program followed: 37° C, 15 minutes; 95° C, 2 minutes; and a 4° C hold.

### Extension

The Extension Thermocycler Program was loaded on the thermocycler and paused at the first step to pre-heat to 98° C until all samples were loaded. Forty-seven (47) μL of the Adaptase Reaction was added, using reagents in the order listed in the manufacturers protocol. The sample was mixed by pipetting or gentle vortexing until homogenous and spun down. The samples were placed in the thermocycler and the following program was run, with lid heating ON. The thermocycler Program followed: 98° C, 30 seconds; 63° C, 15 seconds; 68° C, 5 minutes; and a 4° C hold. Each sample was transferred to a 1.5 mL tube and clean up the Extension Reaction using beads and freshly prepared 80% ethanol was completed.

### Ligation

Twenty (20) μL of the pre-mixed Ligation Reaction Mix was placed in a new PCR tube containing 20 μL of the Post-Extension eluate. Samples were mixed by pipetting or gently vortexing until homogenous and spun down. The samples were placed in the thermocycler programmed at 25 ° C for 15 minutes with lid heating ON, followed by a 4° C hold. Each sample was transferred to a 1.5 mL tube and clean up the Ligation Reaction using beads and freshly prepared 80% ethanol was completed.

### Indexing PCR

Five (5) μL of the appropriate indexed adapter primer(s) were added directly to each sample. Twenty-five (25) μL of the already pre-mixed Indexing PCR Reaction Mix were added to each PCR tube containing 25 μL of sample, using reagents in the order listed in the manufacturers protocol. Samples were mixed by pipetting or gently vortexing until homogenous and spun down. The samples were placed in the thermocycler and the following program run with the proper recommended PCR cycles, with lid heating ON. The thermocycler Program followed: 98° C, 30 seconds; PCR Cycles: 98° C, 10 seconds; 60° C, 30 seconds; 68° C, 60 seconds; 4° C hold. Each sample was transferred to a 1.5 mL tube and clean-up of the Indexing PCR Reaction using beads and freshly prepared 80% ethanol was completed.

### Size Selection/Clean-up

The following protocol was used for each clean-up step, substituting the correct Sample Volume, Bead Volume, and Elution Volume based on the table provided for each section. The magnetic beads were at room temperature and vortexed the beads to homogenize the suspension before use. Each Sample Volume was transferred to a 1.5 mL tube. The specified Bead Volume was added to each sample, mixed by vortex, and quickly spun on a tabletop microcentrifuge. The samples were incubated for 5 minutes at room temperature (off the magnet) and placed on a magnet rack until the solution cleared and a pellet formed (∼2 minutes). The supernatant was removed and discarded without disturbing the pellet (less than 5 μL may have been left behind). A freshly prepared 80% ethanol solution (500 μL) was added to the samples while still on the magnetic rack. Using care not to disturb the pellet, the samples were incubated for 30 seconds, and then the ethanol solution was removed. This step was repeated once more for a second wash with the 80% ethanol solution. The samples were spun in a tabletop microcentrifuge and placed back on the magnetic rack. Residual ethanol solution was removed from the bottom of the tube. The specified Elution Volume of Low TE buffer was added and the pellet re-suspended. Samples were mixed by pipetting up and down until homogenous. Incubation of the samples was completed at room temperature for 2 minutes off the magnet, then placed on the magnet. The entire eluate was transferred to a new 0.2 ml PCR tube. The eluate, without containing the magnetic beads (indicated by brown coloration in the eluate), was ensured to be pure by pipetting the samples into a new tube, placing on a magnet, and transferring the eluate again.

### Microbiome DNA Isolation

Microbiome DNA was collected from blood samples using a modified Microbiome DNA Isolation Kit from NORGEN BIOTEK CORP. (Thorold, ON, Canada). The procedures are as follows:

#### ***A.*** For Samples Collected using Norgen’s Preservation Devices

Transferal of 0.5 mL of whole blood sample to a 2 mL DNAase-free microcentrifuge tube were completed. Lysis Buffer was added and the tube vortexed. One hundred (100) μL of Lysis Additive was added to the mixture and vortexed briefly. The mixture was incubated at 65° C for 5 minutes. The tubes were centrifuged for 2 minutes at 20,000 x g (∼14,000 RPM). The supernatant was transferred to a DNAase-free microcentrifuge tube. One hundred (100) μL of Binding Buffer was added and mixed before incubation on ice. Centrifugation was completed for 2 minutes at 20,000 x g (∼ 14,000 RPM). A pipette was used to transfer up to 700 μL of supernatant into a 2mL DNAase-free microcentrifuge tube. An equal volume of 70% ethanol was added to the lysate collected above and vortexed.

#### ***B.*** Binding to Column

A spin column with one of the provided collection tubes was assembled. Seven hundred (700) μL of the clarified lysate with ethanol was added onto the column and centrifuged for 1 minute at 10,000 x g (∼ 10,000 RPM). The flowthrough was discarded, and the spin column reassembled with the collection tube. The step with the remaining volume of lysate mixture was repeated.

#### ***C.*** Column Wash

Five hundred (500) μL of Binding Buffer was added to the column and centrifuged for 1 minute at 10,000 x g (∼ 10,000 RPM). The flowthrough was discarded, and the spin column reassembled with its collection tube. Five hundred (500) μL of Wash Solution was applied to the column and centrifuged for 1 minute at 10,000 x g (∼ 10,000 RPM). The flow-through was discarded and the spin column reassembled with its collection tube. Repeat the previous two steps. The column was centrifuged for 2 minutes at 20,000 x g (∼ 14,000 RPM) in order to thoroughly dry the resin. The collection tube was discarded.

#### ***D.*** DNA Elution

The column was placed into a fresh 1.7 mL Elution tube provided with the kit. Fifty (50) μL of Elution Buffer was added to the column. Centrifugation was completed for 1 minute at 200 x g (∼ 2,000 RPM), followed by a 1-minute spin at 20,000 xg (∼14,000 RPM). An additional elution was performed using 50 μL of the Elution Buffer.

#### ***E.*** Storage of DNA

The purified genomic DNA was stored at −80° C.

### Sequencing

Quality Assurance/Quality Control of bar-coded sequence prepped samples of cDNA were completed by The Genomics Core Facility at Oklahoma State University (Stillwater, OK). The Genomics Core Facility at Oklahoma State University (Stillwater, OK) completed the cDNA library sequencing with an illumina RNASeq NGS instrument.

Quality Assurance/Quality Control of bar-coded sequence prepped samples were completed and sequenced for 16S rRNA metagenomics of blood samples by Swift Biosciences^TM^ (Ann Arbor, MI USA) using an illumina MiSeq NGS instrument.

### Computational Analysis

Several methods were utilized to conduct bioinformatic analysis of the obtained sequence data. For gene expression analyses, we used the Partek^®^ Flow^®^ software suites pipelines that include, but are not limited to the STAR algorithm, Normalization, and the gene set differential analysis method (GSA).

Preliminary analyses included Qiime2 analysis for the metagenomic or microbiome analysis. We present here, the results obtained utilizing the Kraken pipeline through the Partek^®^ Flow^®^ software suites, a start-to-finish software analysis solution for next generation sequencing data applications. Inflammation comparisons were based on the ratio of *Firmicute*s/*Bacteroidetes* (*F*/*B* Ratios) that are evident as well as the reduction of microbial diversity.

## Results

We analyzed gene expression in samples from 19/20 wethers (7 dpi). The microbiome portion resulted in analysis of 38/40 wethers separated into 7 dpi (18/20 samples) and 21 dpi (20/20 samples). Initially, ten (10) individuals were not infected with *H. contortus*, 10 received infection with *H. contortus* only, 10 were infected with *H. contortus* and received ZA injections, and 10 were infected with *H. contortus* and received AB injection. As mentioned in this study blood samples were collected 7 dpi, where 19/20 cDNA libraries passed QA/QC analytics for cDNA gene expression analysis. QA/QC analytics of high-throughput barcoded illumina MiSeq NGS sequencing for 16S rRNA metagenomics, resulted in the 18/20 samples being obtained for 7 dpi and the 20/20 samples for 21 dpi. The sequences used for the purposes of this study, involved the inclusion of gene expression and metagenomic analyses. The initial design for the study is shown for 20 individuals in Table 2 (Supplementary Documents).

**Table 2.**
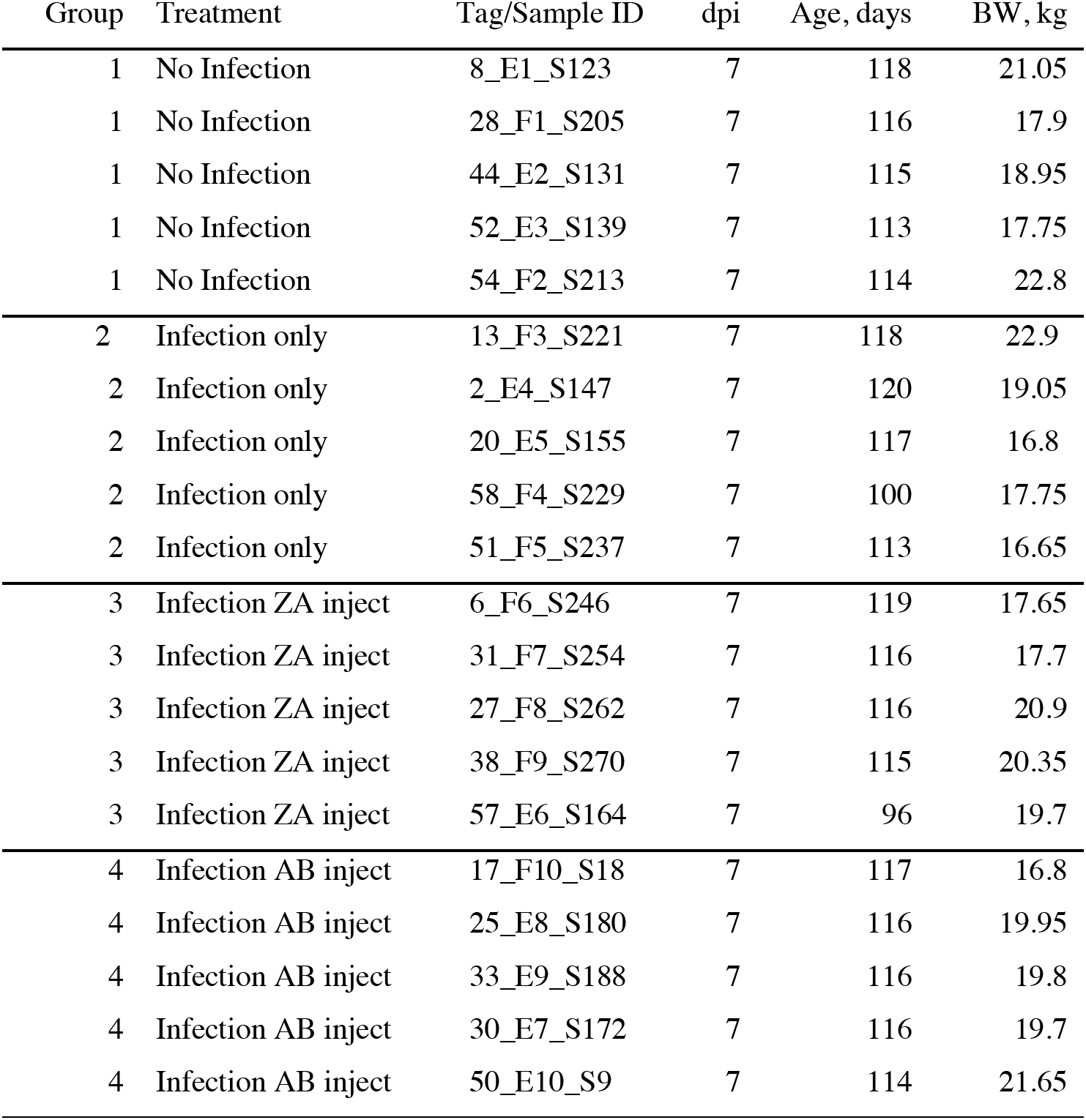
Detailed Experimental Set-up. The Group number, Treatment type, identifying Tag, days post inoculation (dpi), Age in days, and Body Weight (BW) in kg.

*Haemonchus contortus* infection affected the metabolic system of wethers. A 95% confidence interval (* = p < 0.05) indicated likely significant changes in the expression of at least 184 genes (affecting treatment comparisons of samples that were infected with zoledronic acid injection versus infection with antibody injection Fig. 7) when using a numeric triad of p-values for no infection versus infection only, no infection versus infection zoledronic acid injection, and no infection versus infection with antibody injection. The hierarchical clustering/heat map (Figs. 1) generated after selection of three of the most highly significant specific differentially analyzed genes, depicts colored tiles showing differences in genomic features in the integration site data sets from the blood samples of each subject (n = 19). They indicate the intensity and direction of any significant departures from the distributions of random controls varying to a degree depending on the type of treatment and the subject [19].

**Fig. 1.**
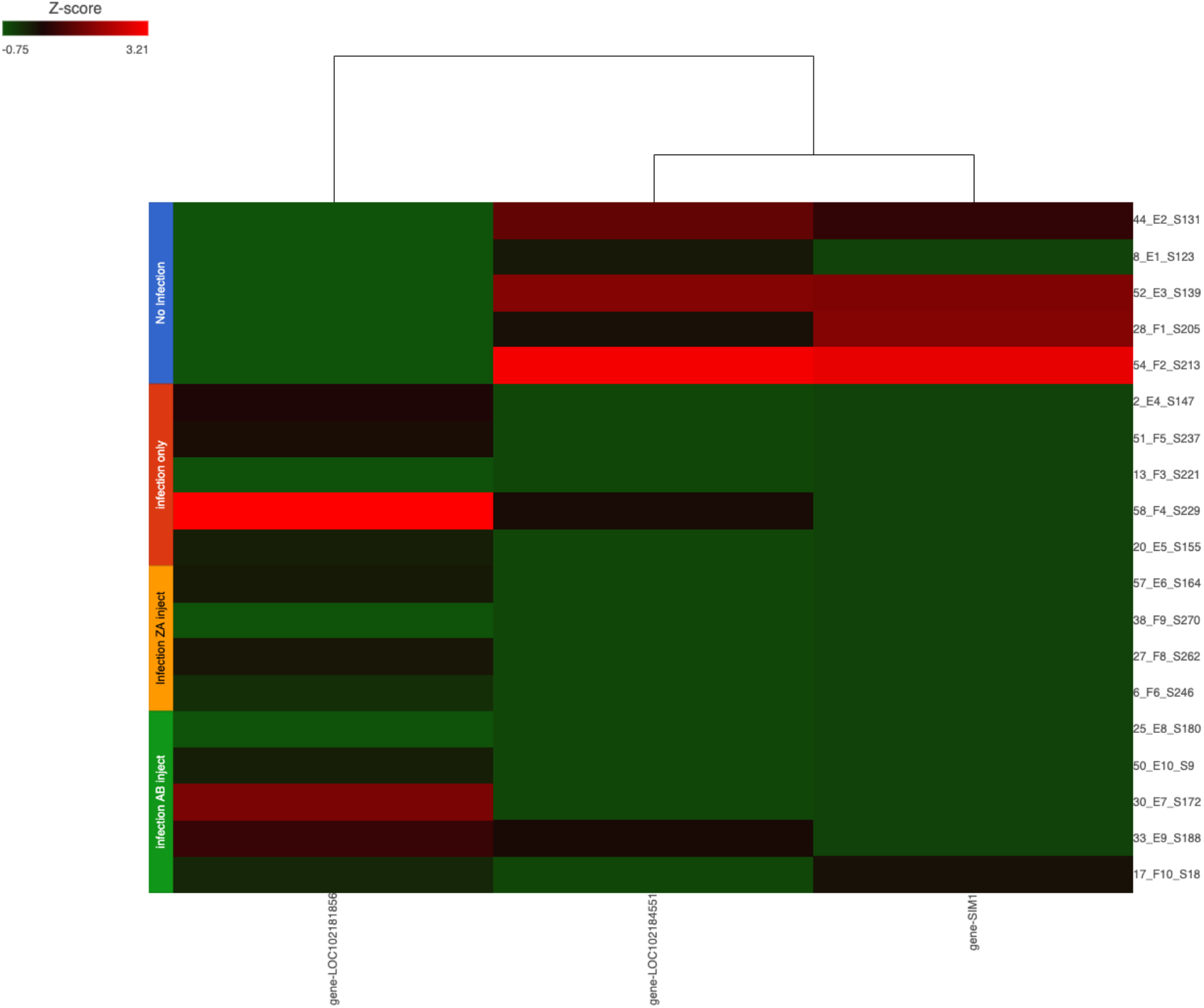
Hierarchical clustering/heat map. Three selected genes (identified at the bottom of the image) following differential analysis using GSA counts of 19/20 samples (identified on the right side of the image belonging to the described treatment group identified on the left side of the image) indicate the most likely significant genes (p < 0.05) affecting samples when comparing No Infection samples to those that were only infected with the *H. contortus* pathogen, injected with zoledronic acid (ZA) or injected with antibodies (AB) and then infected with the *H. contortus* pathogen on 7 dpi.

The following Volcano plots (Figs. 2–7) show genes that were identified as being downregulated (* = < −2), having no fold change (NC: * = > −2, * = < 2), and/or being upregulated (* = > 2). The Volcano plots illustrates the related distribution of genes out of 7627 expressed, depending on subject and treatment type. The distribution of genes in this case were based on comparisons using a numeric triad of p-values for no infection versus infection only, no infection versus infection zoledronic acid (ZA) injection, and no infection versus infection with antibody (AB) injection. Significance was based on a 95% confidence interval (* = p < 0.05) where fold changes of downregulated, NC, and upregulated gene distributions of treatment groups were assessed. In Fig. 2 a comparison of no infection versus infection only subjects, out of 7627 genes that were expressed, 523 genes were significant in expression during downregulation, NC, and upregulation. For the comparison of no infection vs infection ZA (Fig. 3), 290 genes that were significant in expression during downregulation, NC, and upregulation. In Fig. 4 289 genes were significant in expression during downregulation, NC, and upregulation when comparing no infection vs infection AB. In Fig. 5 examines the comparison of infection only vs infection ZA, resulting in the identification of 338 genes that were expressed with significance in expression during downregulation, NC, and upregulation. The Volcano plot for Fig. 6 indicates 275 genes that were expressed in fold changes for downregulated, NC, and upregulated genes that were significant.

**Fig. 2.**
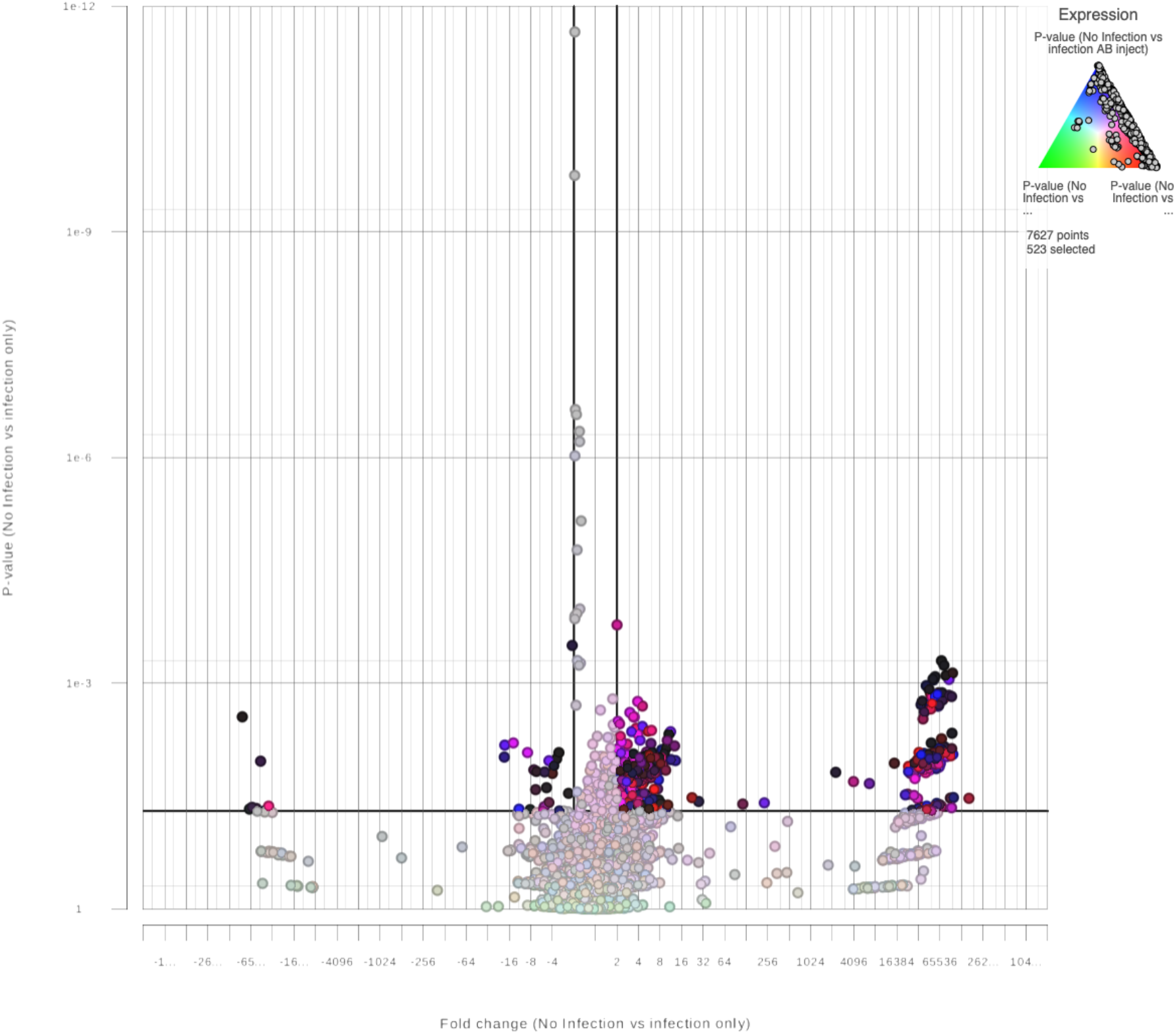
No Infection vs Infection only. A Volcano plot for 19/20 blood samples indicating likely significant expression of genes (* = p < 0.05) based on treatment type of *Capra hircus* following STAR alignment and GSA differential analysis for transcript sequences of 7627 identified genes expressed on 7 dpi. The fold change indicates downregulated (* = < −2), no change (NC; * = > −2, * = < 2), and upregulated (* = > 2) gene distribution when comparing No Infection samples to infection only samples when expressed against a Numeric Triad of P-values for No Infection vs Infection only (green), No Infection vs Infection ZA inject (red) and No Infection vs infection AB inject (blue).

**Fig. 3.**
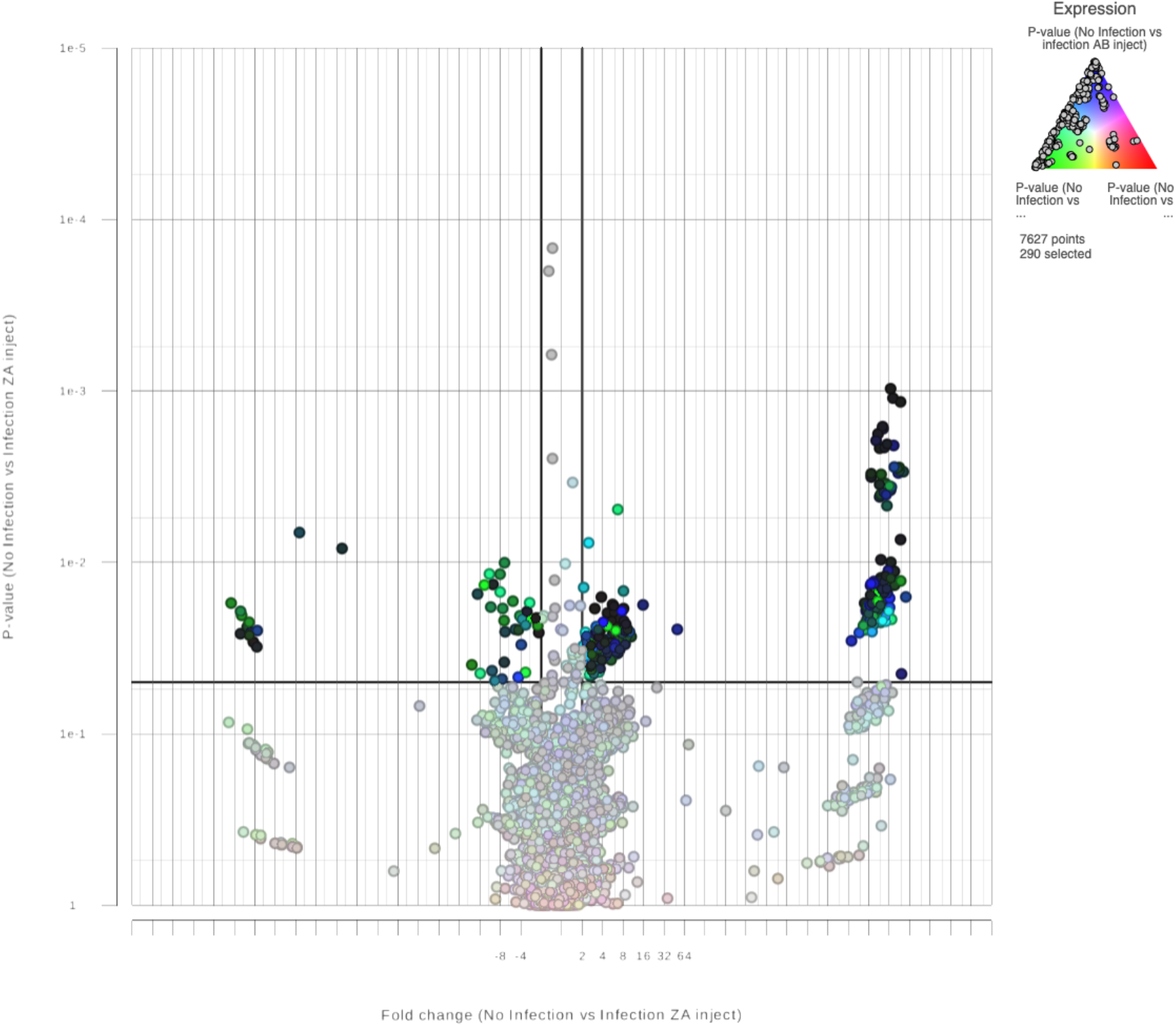
No Infection vs Infection ZA inject. A Volcano plot for 19/20 blood samples indicating likely significant expression of genes (* = p < 0.05) based on treatment type of *Capra hircus* following STAR alignment and GSA differential analysis for transcript sequences of 7627 identified genes expressed on 7 dpi. The fold change indicates downregulated (* = < −2), no change (NC; * = > −2, * = < 2), and upregulated (* = > 2) gene distribution when comparing No Infection samples to Infection ZA inject samples when expressed against a Numeric Triad of P-values for No Infection vs Infection only (green), No Infection vs Infection ZA inject (red) and No Infection vs infection AB inject (blue).

**Fig. 4.**
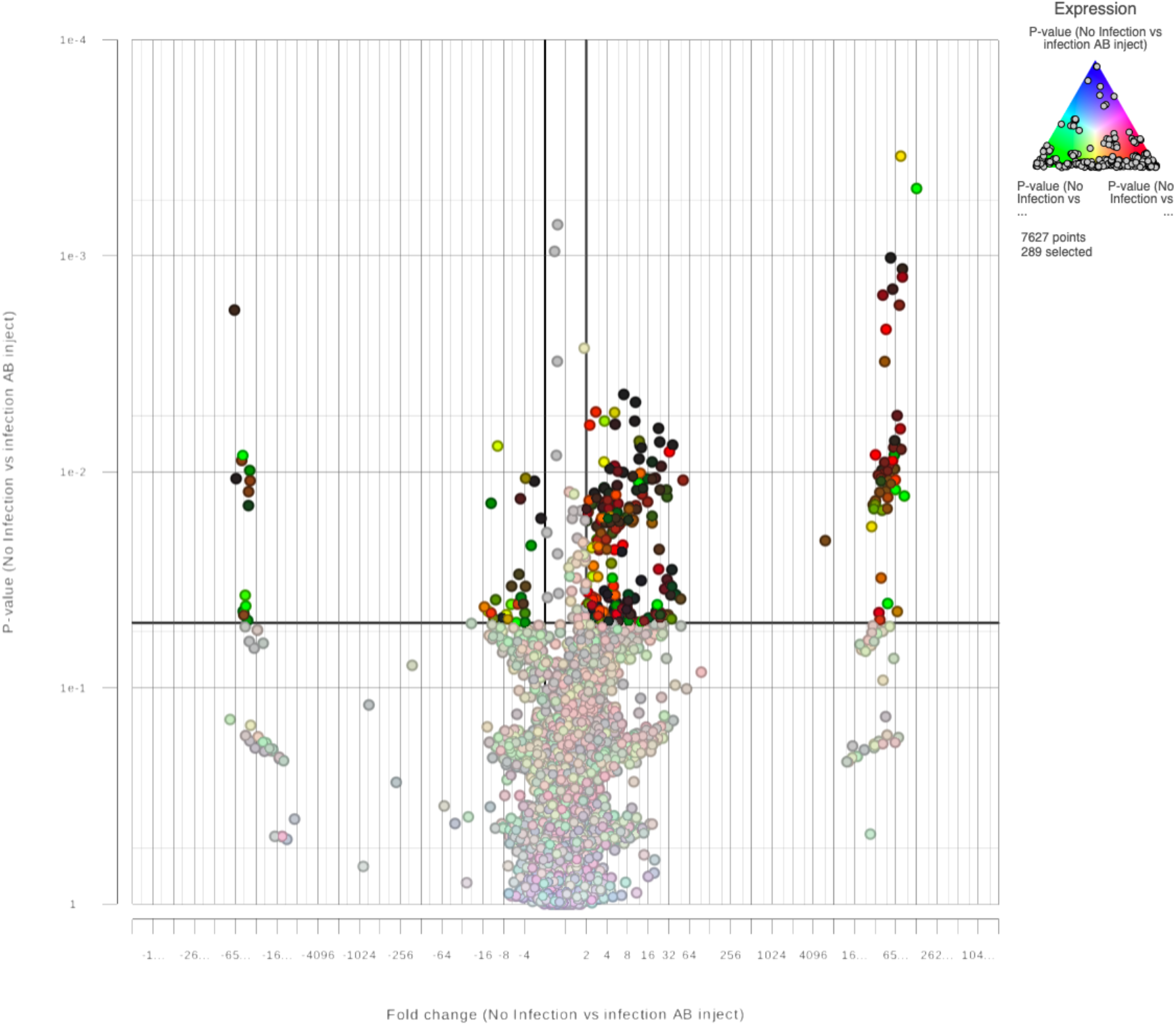
No Infection vs Infection AB inject. A Volcano plot for 19/20 blood samples indicating likely significant expression of genes (* = p < 0.05) based on treatment type of *Capra hircus* following STAR alignment and GSA differential analysis for transcript sequences of 7627 identified genes expressed on 7 dpi. The fold change indicates downregulated (* = < −2), no change (NC; * = > −2, * = < 2), and upregulated (* = > 2) gene distribution when comparing No Infection samples to infection AB inject samples when expressed against a Numeric Triad of P-values for No Infection vs Infection only (green), No Infection vs Infection ZA inject (red) and No Infection vs infection AB inject (blue).

**Fig. 5.**
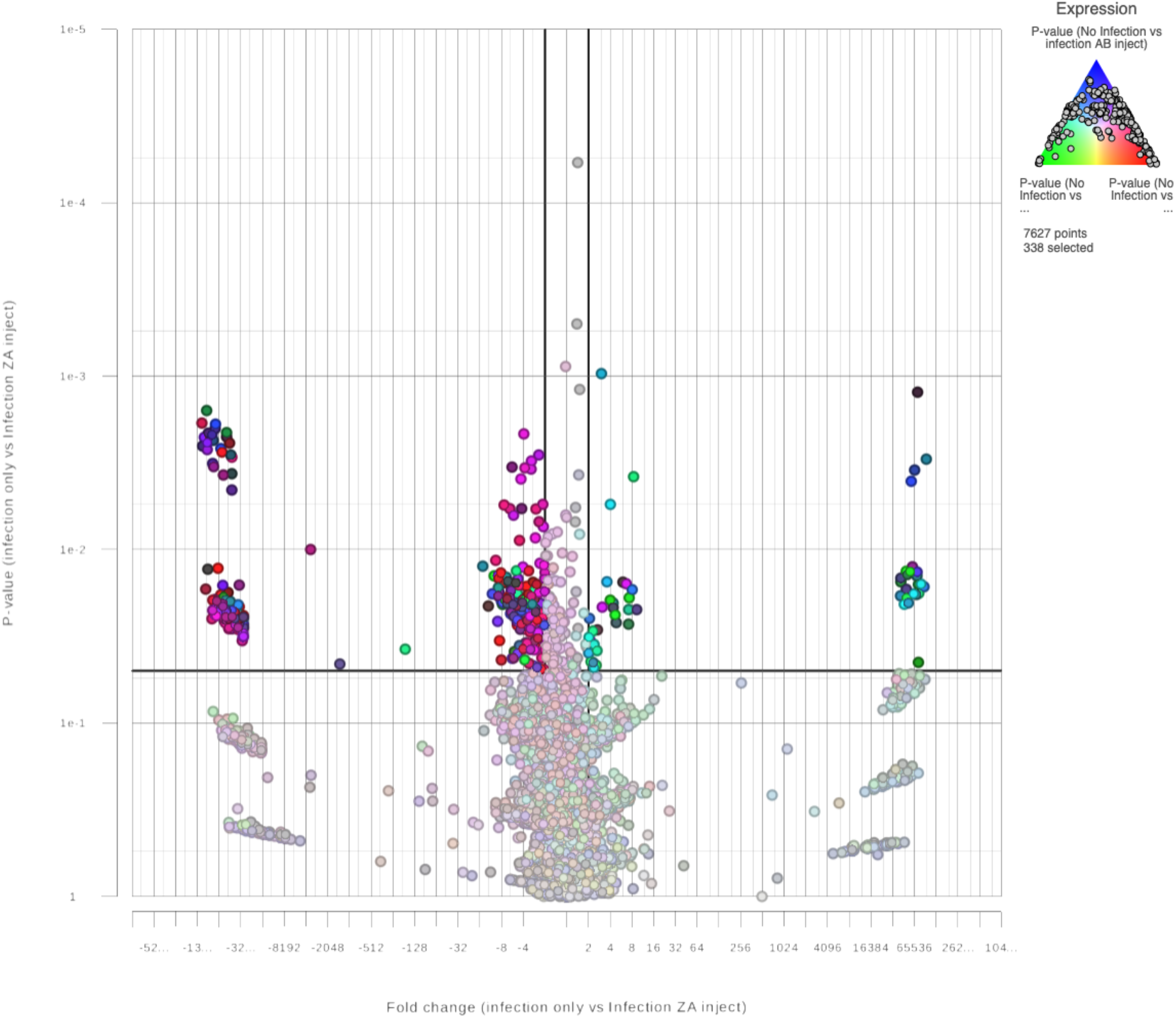
Infection only vs Infection ZA inject. A Volcano plot for 19/20 blood samples indicating likely significant expression of genes (* = p < 0.05) based on treatment type of *Capra hircus* following STAR alignment and GSA differential analysis for transcript sequences of 7627 identified genes expressed on 7 dpi. The fold change indicates downregulated (* = < −2), no change (NC; * = > −2, * = < 2), and upregulated (* = > 2) gene distribution when comparing infection only samples to Infection ZA inject samples when expressed against a Numeric Triad of P-values for No Infection vs Infection only (green), No Infection vs Infection ZA inject (red) and No Infection vs infection AB inject (blue).

**Fig. 6.**
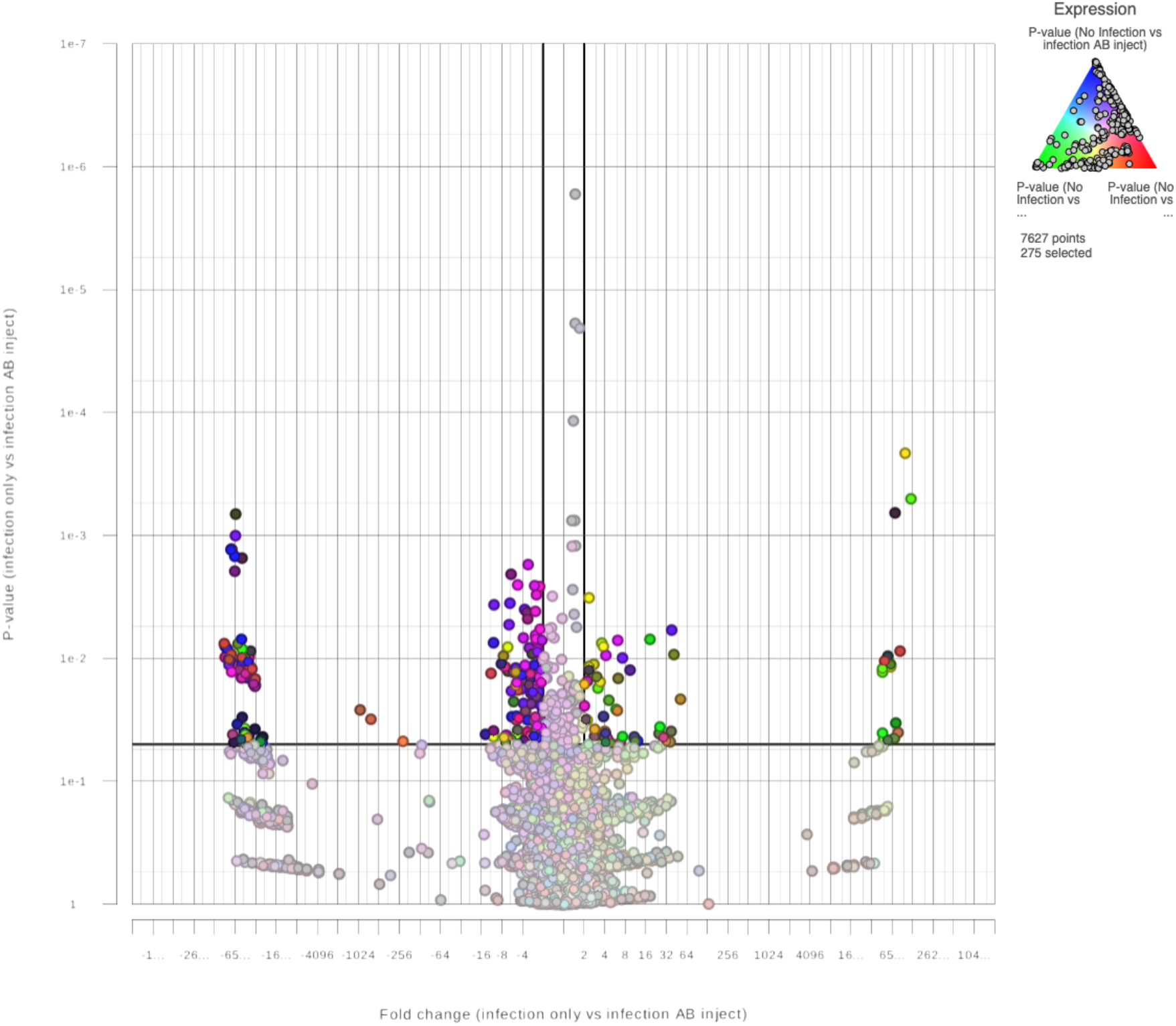
Infection only vs Infection AB inject. A Volcano plot for 19/20 blood samples indicating likely significant expression of genes (* = p < 0.05) based on treatment type of *Capra hircus* following STAR alignment and GSA differential analysis for transcript sequences of 7627 identified genes expressed on 7 dpi. The fold change indicates downregulated (* = < −2), no change (NC; * = > −2, * = < 2), and upregulated (* = > 2) gene distribution when comparing infection only samples to infection AB inject samples when expressed against a Numeric Triad of P-values for No Infection vs Infection only (green), No Infection vs Infection ZA inject (red) and No Infection vs infection AB inject (blue).

**Fig. 7.**
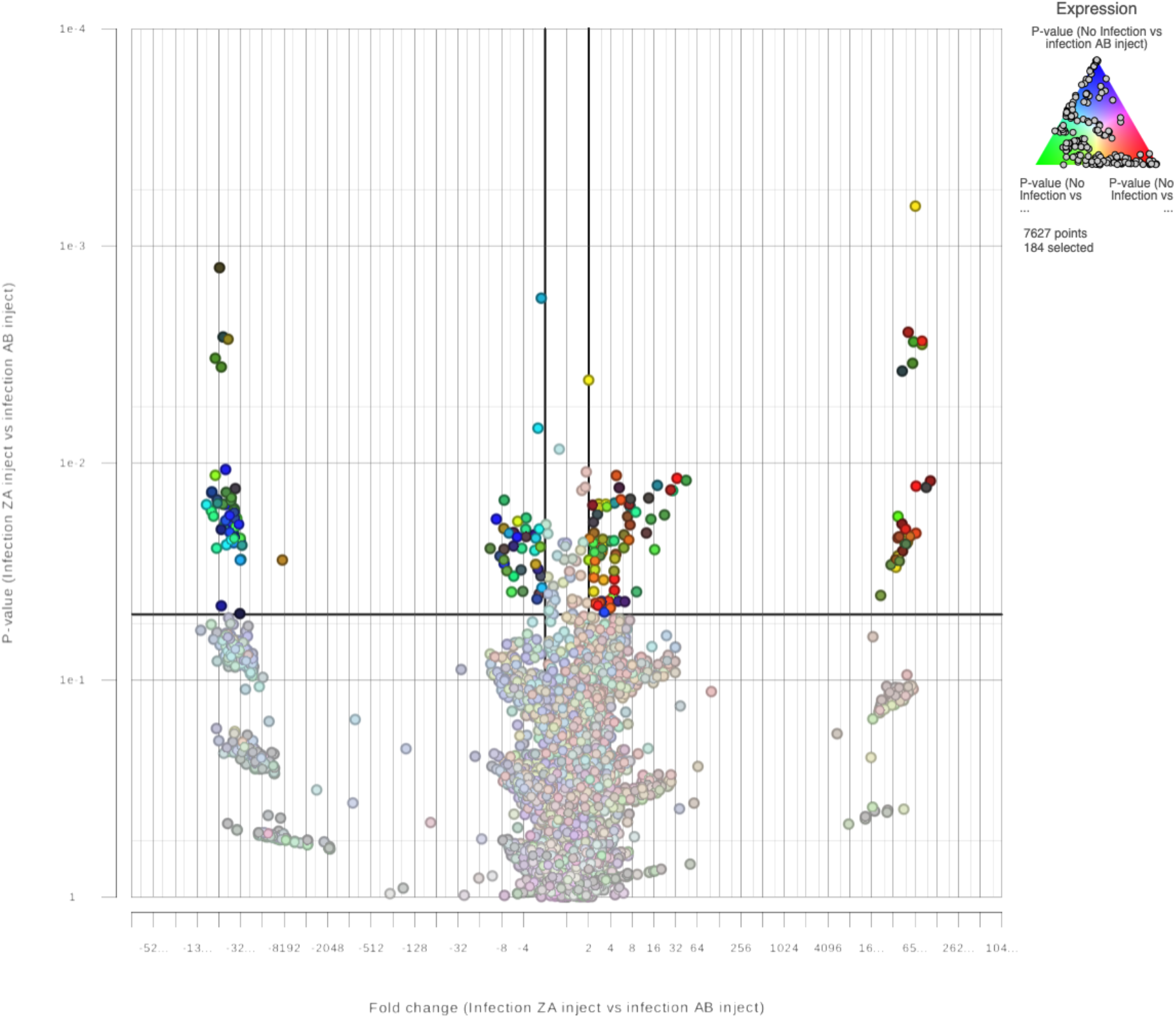
Infection ZA inject vs Infection AB inject. A Volcano plot for 19/20 blood samples indicating likely significant expression of genes (* = p < 0.05) based on treatment type of *Capra hircus* following STAR alignment and GSA differential analysis for transcript sequences of 7627 identified genes expressed on 7 dpi. The fold change indicates downregulated (* = < −2), no change (NC; * = > −2, * = < 2), and upregulated (* = > 2) gene distribution when comparing Infection ZA inject samples to infection AB inject samples when expressed against a Numeric Triad of P-values for No Infection vs Infection only (green), No Infection vs Infection ZA inject (red) and No Infection vs infection AB inject (blue).

The average number of raw reads for metagenomic analyses of blood samples using Kraken is illustrated in Figs. 8 and 9. The distribution of raw read counts indicate the apparent difference in number between 7 dpi and 21 dpi subjects. There were 43.61 raw reads for metagenomic analysis of blood collected on 7 dpi as compared to 2,638.80 raw reads for metagenomic analysis of blood samples at 21 dpi.

**Fig. 8.**
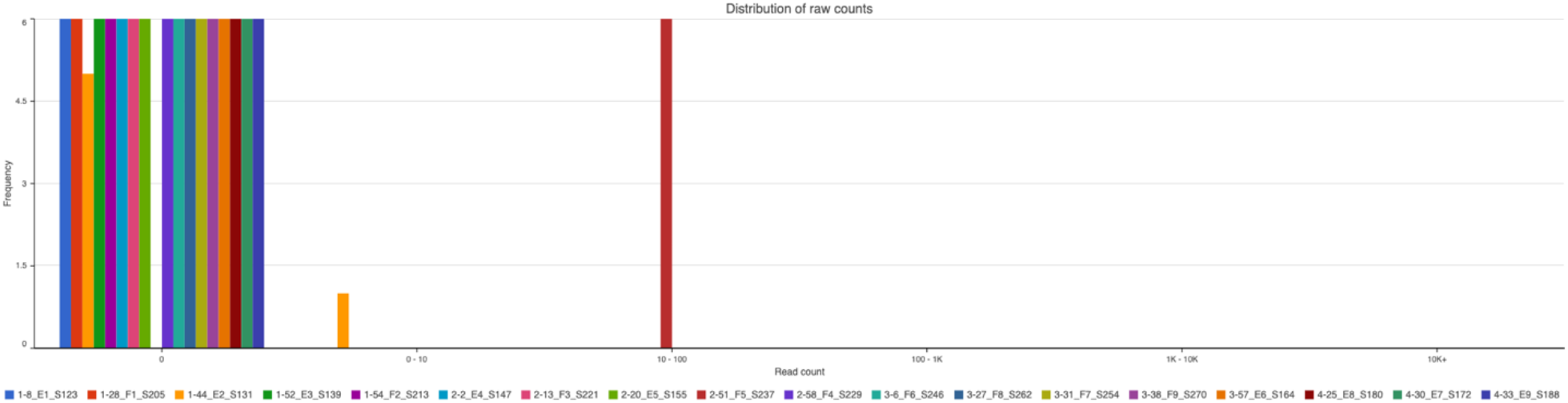
Distribution of raw read counts for all samples at 7 dpi. The first number before each sample ID identifies the treatment type (1 = No infection, 2 = Infection only, 3 = Infection ZA inject, and 4 = Infection AB inject).

**Fig. 9.**
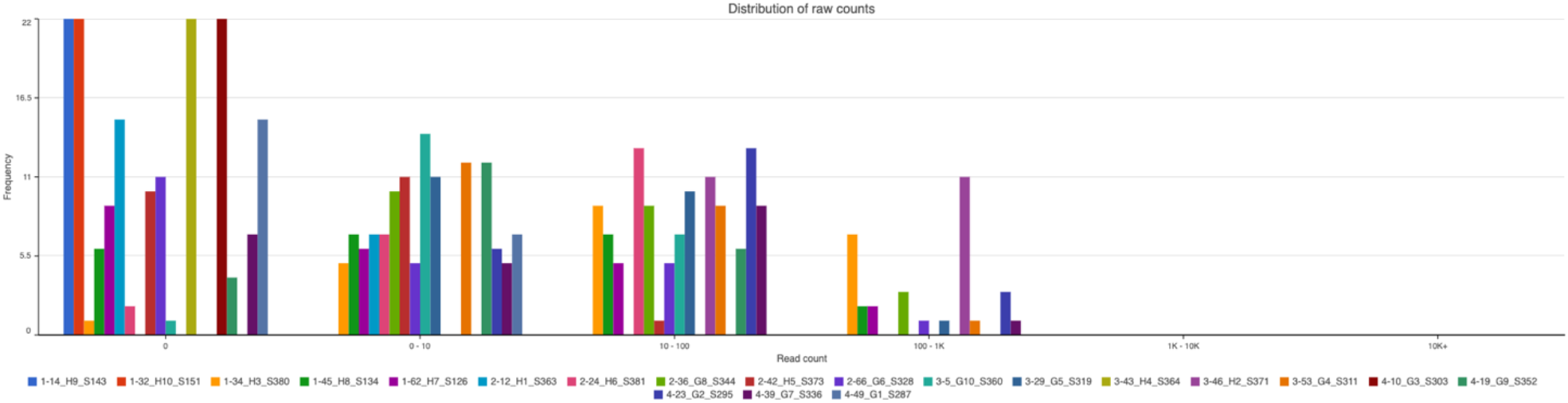
Distribution of raw read counts for all samples at 21 dpi. The first number before each sample ID identifies the treatment type (1 = No infection, 2 = Infection only, 3 = Infection ZA inject, and 4 = Infection AB inject).

After QA/QC, we collected Alpha diversity reports for 18 individuals for 7 dpi using Shannon and Simpson distribution indices (Fig. 10) [12]. The Alpha diversity reports for the Shannon and Simpson distribution indices (Table 3; Supplementary Documents) were evaluated. Our results indicate that *H. contortus* has a significant effect on species-level microbial diversity for blood that is infected and injected with ZA using both indices [P(T<=t) first-tail 0.012 and P(T<=t) second-tail 0.023] (Table 4).

**Fig. 10.**
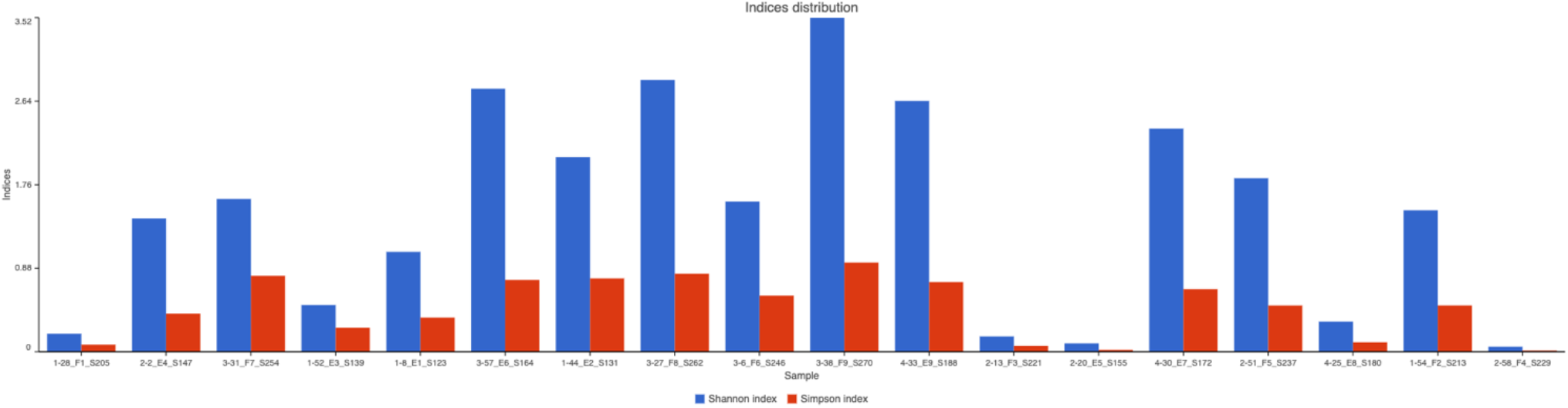
Shannon and Simpson indices of 7dpi. **Alpha** diversity reports depicted by Shannon and Simpson indices distributions for 18 samples of 7dpi (loss of sample numbers 17 and 50 due to poor quality) [23–25] where single samples and the variation of microbes in them are identified. The first number before each sample ID identifying the treatment type (1 = No infection, 2 = Infection only, 3 = Infection ZA inject, and 4 = Infection AB inject).

**Table 3.**
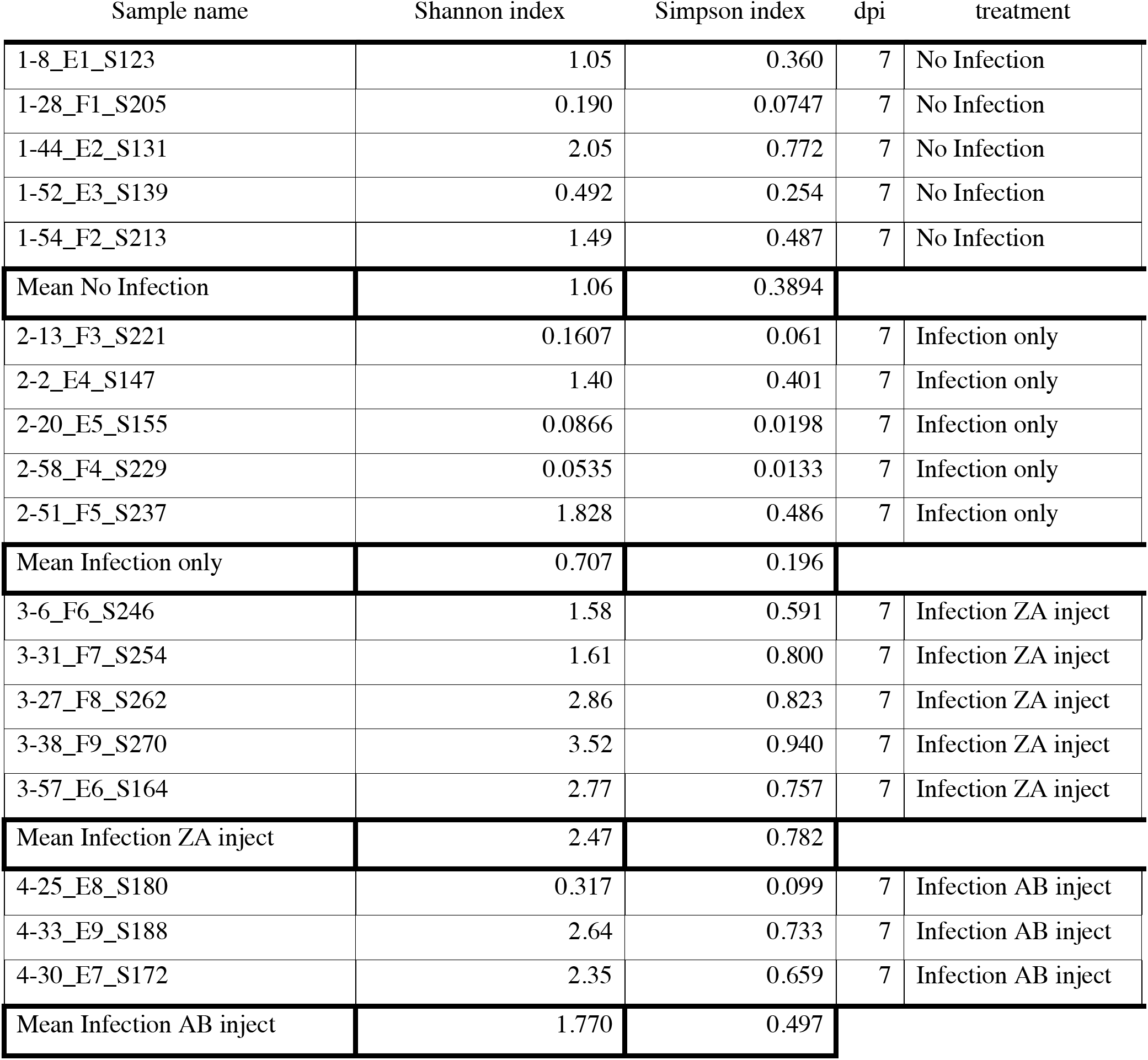
Alpha diversity report for 7 dpi Shannon and Simpson index.

**Table 4.**
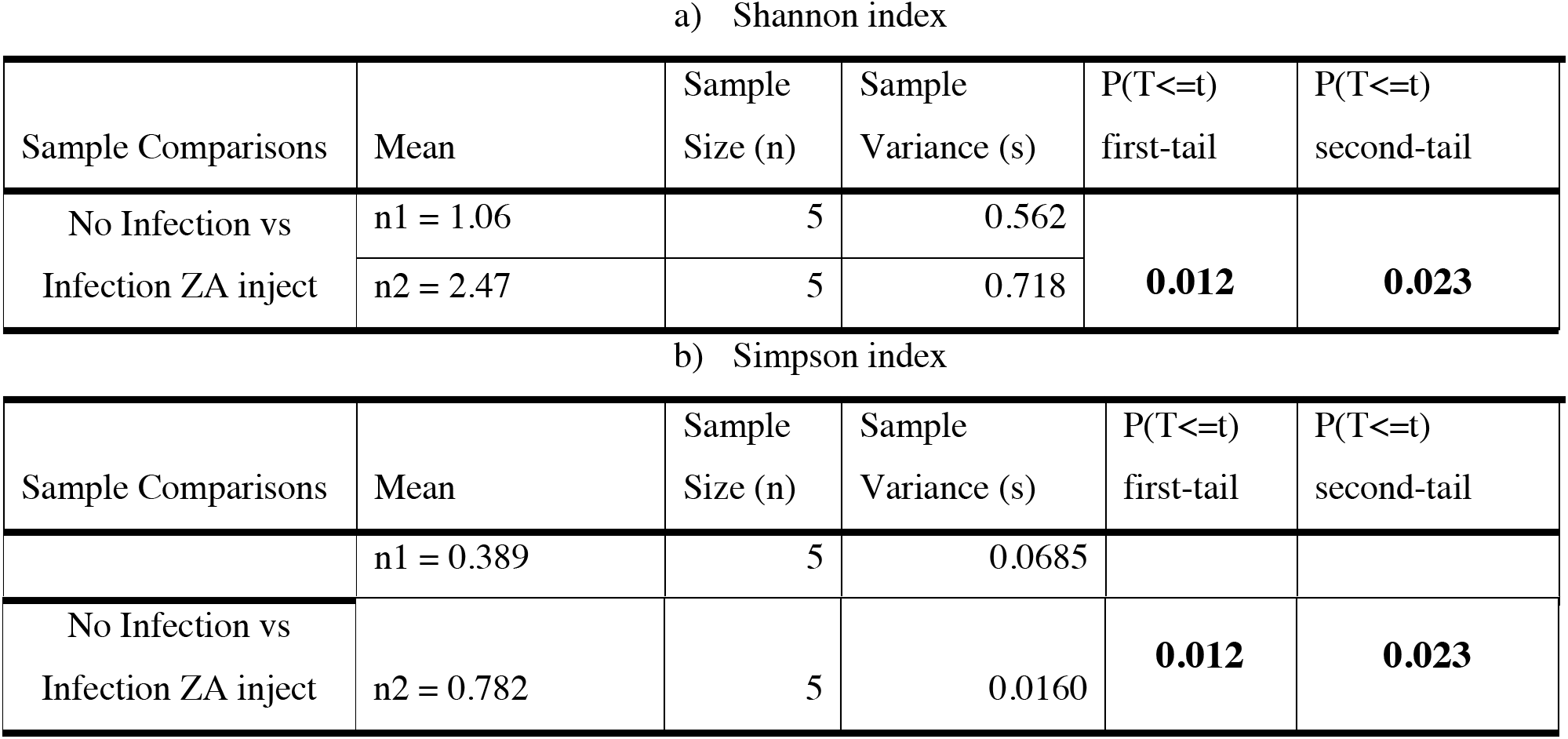
Significant values of 7 dpi Shannon and Simpson indices. Statistical comparison of a) Shannon and b) Simpson distribution indices for Alpha diversity reports of 7 dpi richness and diversity of microbial flora in host *Capra hircus* wethers.

b) Simpson index
| Sample Comparisons | Mean | Sample Size (n) | Sample Variance (s) | P(T<=t) first-tail | P(T<=t) second-tail |
| --- | --- | --- | --- | --- | --- |
|  | n1 = 0.389 | 5 | 0.0685 |  |  |
| No Infection vs<br>Infection ZA inject | n2 = 0.782 | 5 | 0.0160 | **0.012** | **0.023** |

We also collected Alpha diversity reports for 20 individuals for 21 dpi (Table 4; Supplementary Documents; Fig. 11).

**Fig. 11.**
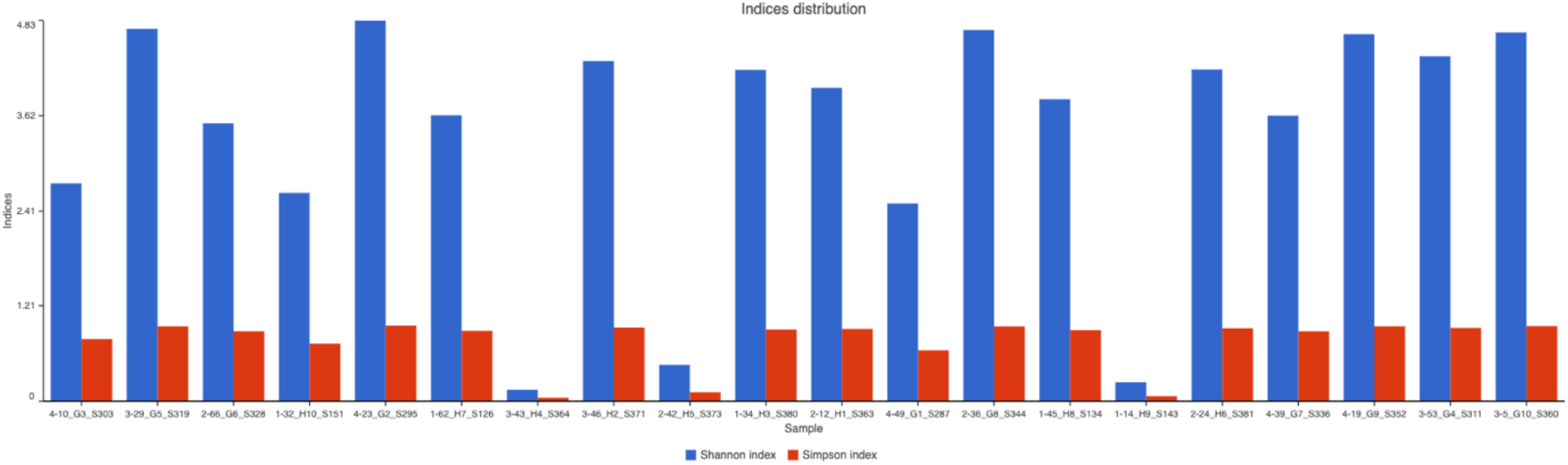
Shannon and Simpson indices of 21dpi. Alpha diversity reports depicted by Shannon and Simpson distribution indices for 20 samples of 21dpi. The first number before each sample ID identifies the treatment type (1 = No infection, 2 = Infection only, 3 = Infection ZA inject, and 4 = Infection AB inject).

Species richness and diversity for non-infected controls, infected wethers, infected ZA injected wethers, and infected AB injected wethers indicated significant differences between 7dpi and 21 dpi microbial composition. This contrasts with 21 dpi where there are no significant differences when comparisons are made between non-infected and infected wethers (ZA/AB injected or not). This supports literature where the total bacterial load increases over time after infection with *H. contortus* [8].

Infection likely has a broad range of quantitative biological effects on each host. Our investigation of the composition of microbial flora shows a more similar pattern to other studies that support the notion as to why a host is better suited to withstand intrusion by external threats. The richer and more diverse the microbiota, the better the host may combat external pathogens [12]. The Alpha diversity report, depicting Shannon and Simpson distribution indices, identifies that treatments for 21 dpi (Table 5; Supplementary Documents), when compared with each other do not show significance. The non-infected wethers appear to have developed a pronounced, non-infected version of an abundant and diverse microbial flora. Thus, when infected 21 dpi wethers are statistically compared to non-infected 21 dpi controls for abundance and diversity using Alpha diversity reports with Shannon and Simpson distribution indices, there are no significant differences.

**Table 5.**
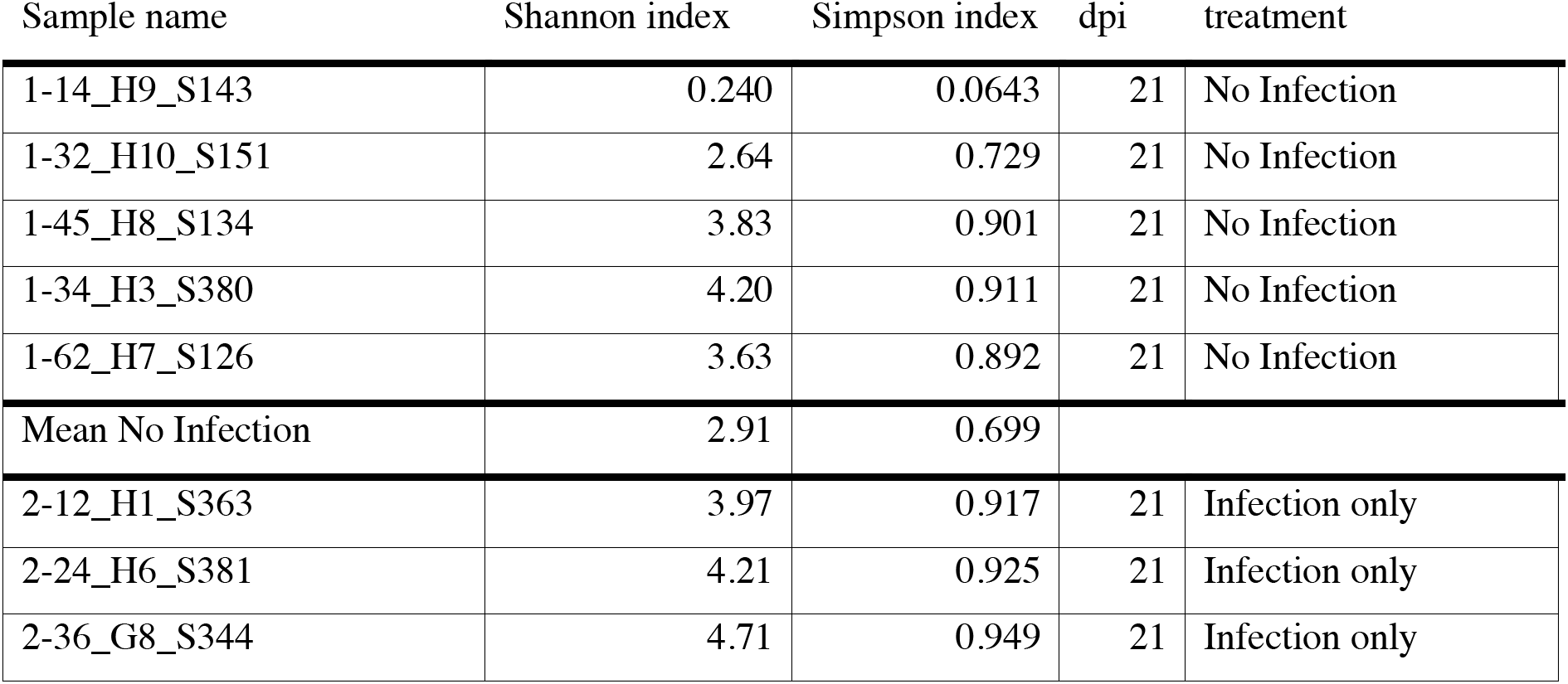
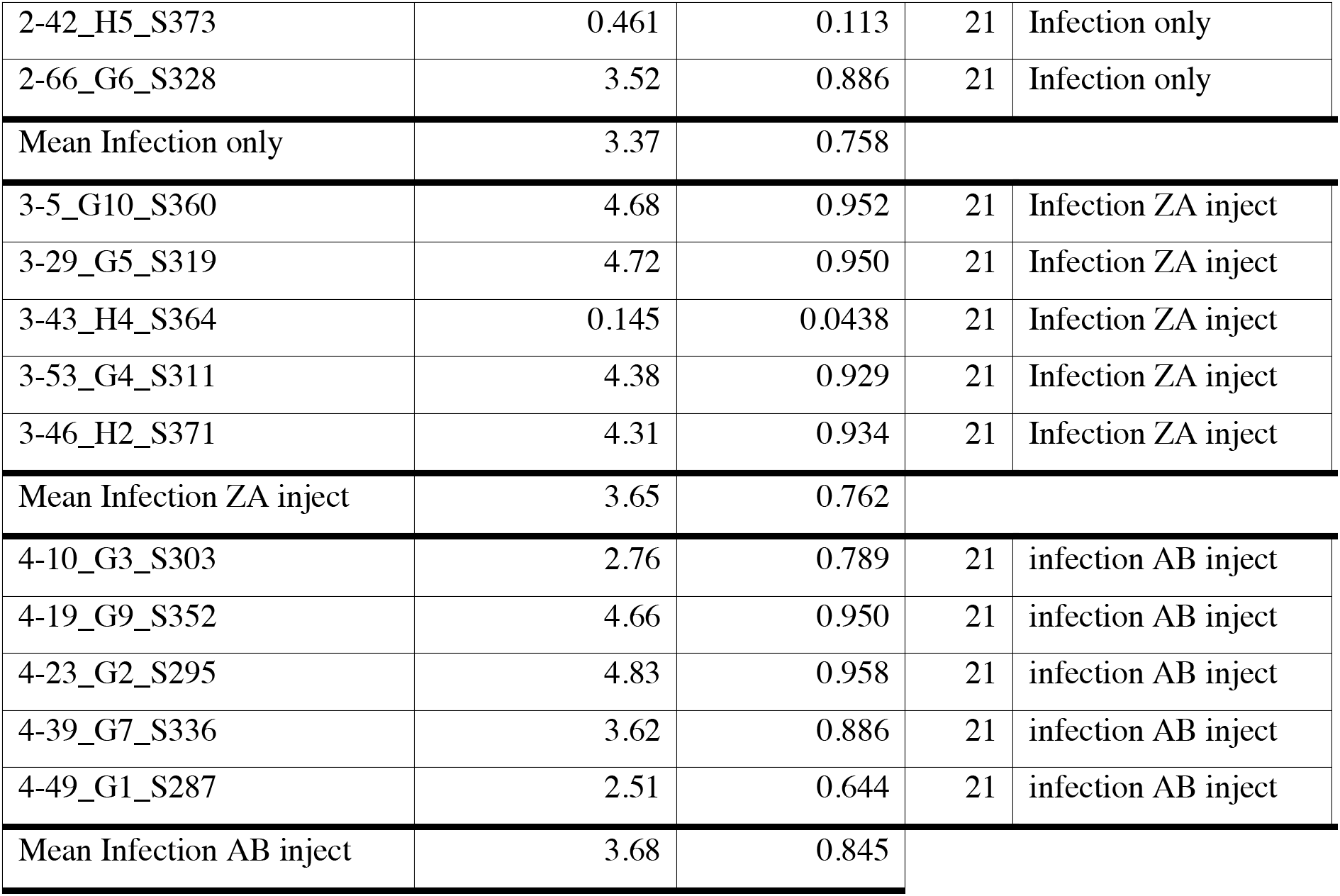
Alpha diversity report for 21 dpi Shannon and Simpson index.

However, when comparing Shannon and Simpson Alpha indices between 7 dpi and 21 dpi (Table 6) there are significant differences. The Shannon indices for 7 dpi vs 21 dpi t-Test: Two-Sample Assuming Unequal Variances results showed that there are significant differences between the different treatment groups. There are differences between “non-infected” wethers based on age alone. Table 5 shows that there are significant differences between comparisons between “non-infected” wethers for 7 dpi and those wethers that were “non-infected” for 21dpi, only “infected” for 21 dpi, “infected” with ZA injection for 21 dpi, and “infected” with an AB injection for 21 dpi. Therefore, it is evident that the richness and diversity for microbial flora in blood changes over time, regardless of the treatment. Further evidence supports this when examining wethers subjected to “infection” only for 7 dpi as compared to those with “non-infected” 21 dpi following inoculation, “infection” with ZA injection for 21dpi, and “infection” with an AB injection for 21 dpi. This is further support that despite the condition, as the age of the wethers increases (dpi), there are significant changes in the richness and diversity of microbial flora in the blood.

**Table 6.**
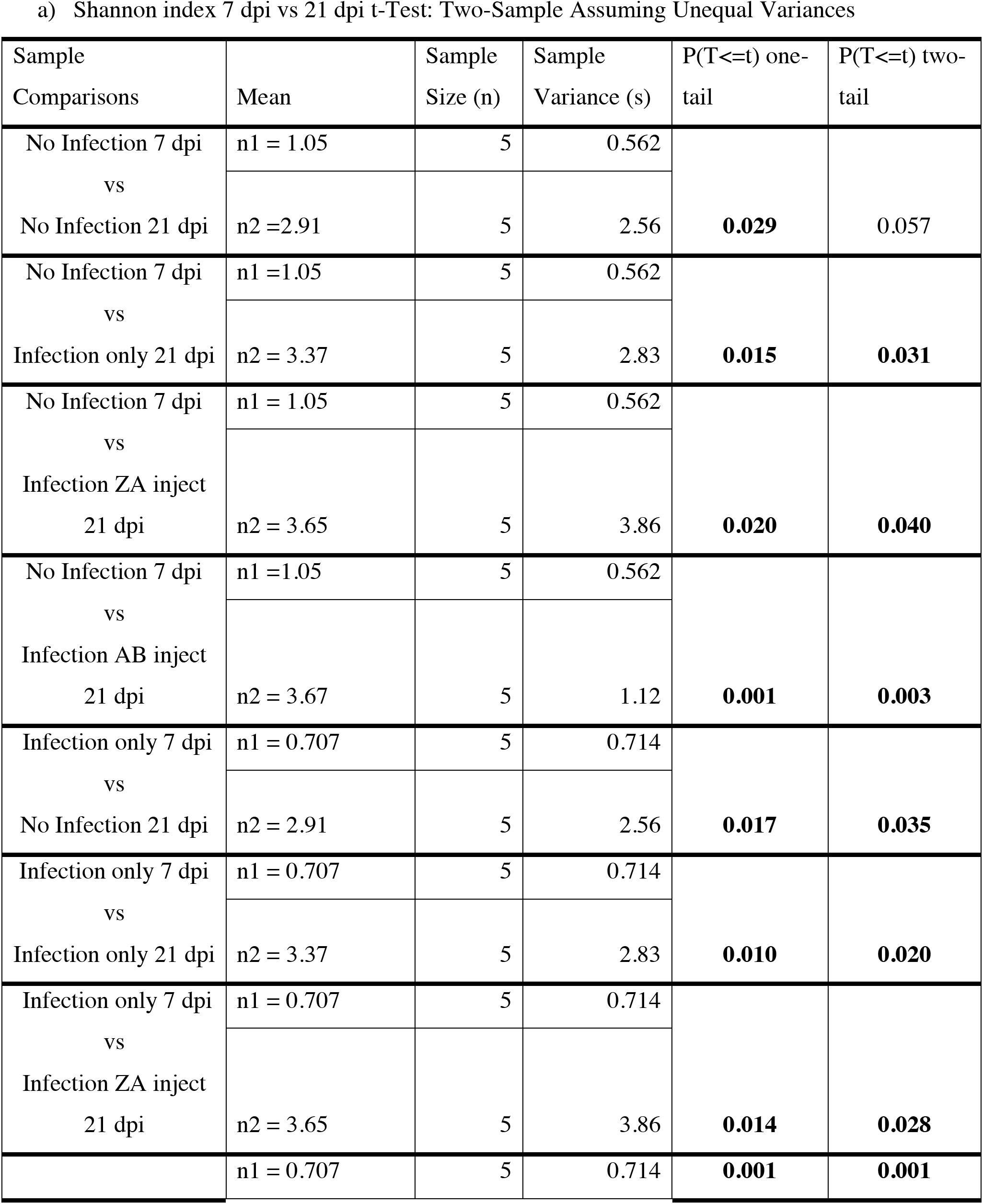
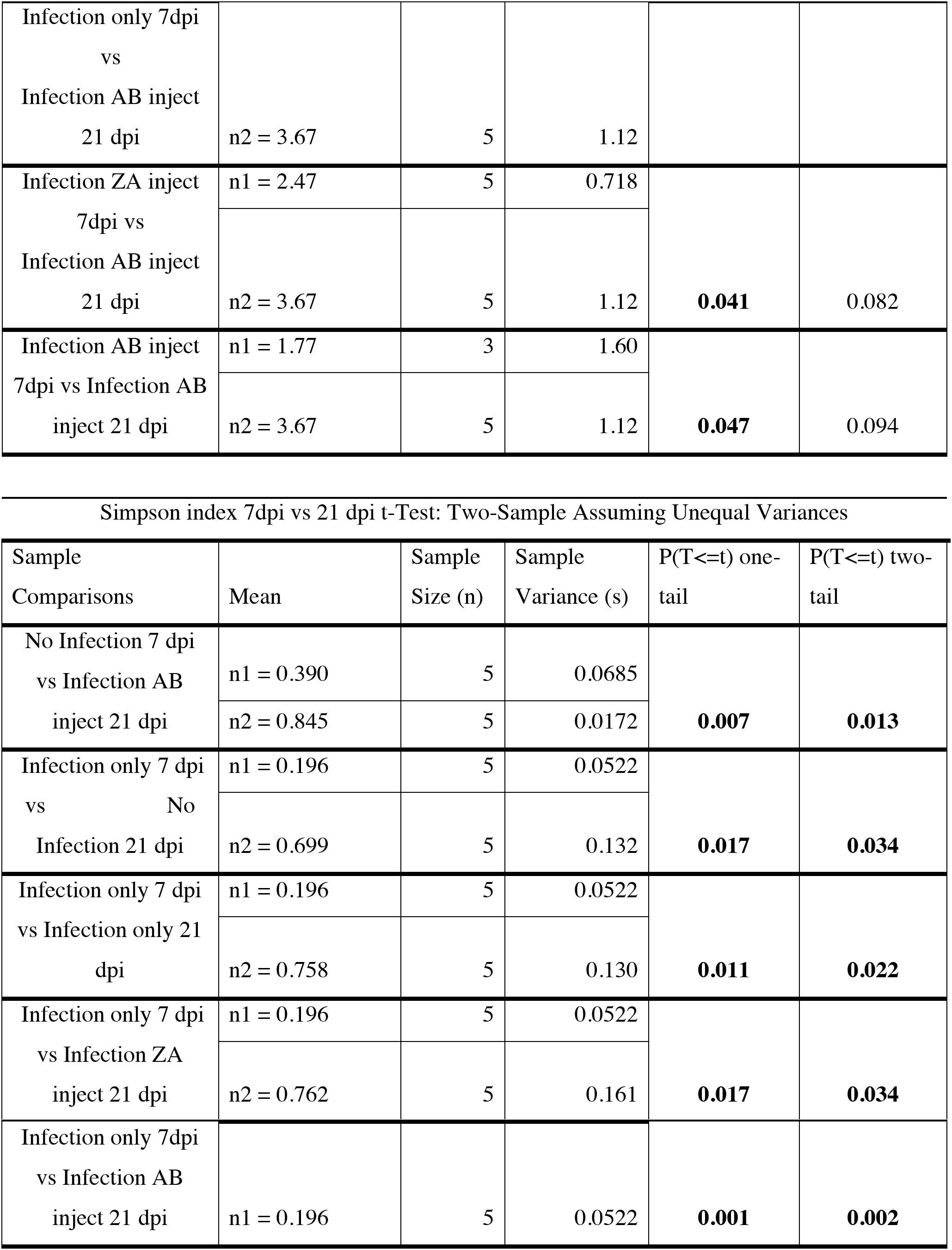
Statistically different comparison results of 7 dpi versus 21 dpi. Statistically different comparison results of 7 dpi versus 21 dpi a) Shannon and b) Simpson distribution indices for Alpha diversity reports of abundance and diversity for microbial flora in host *Capra hircus* wethers.

Gene amplicons for 16S rRNA Relative Abundance profiles are illustrated for 18 individuals on 7dpi (Fig. 12). The blood samples are dominated by 31 most abundant operational taxonomic units (OTUs).

**Fig. 12.**
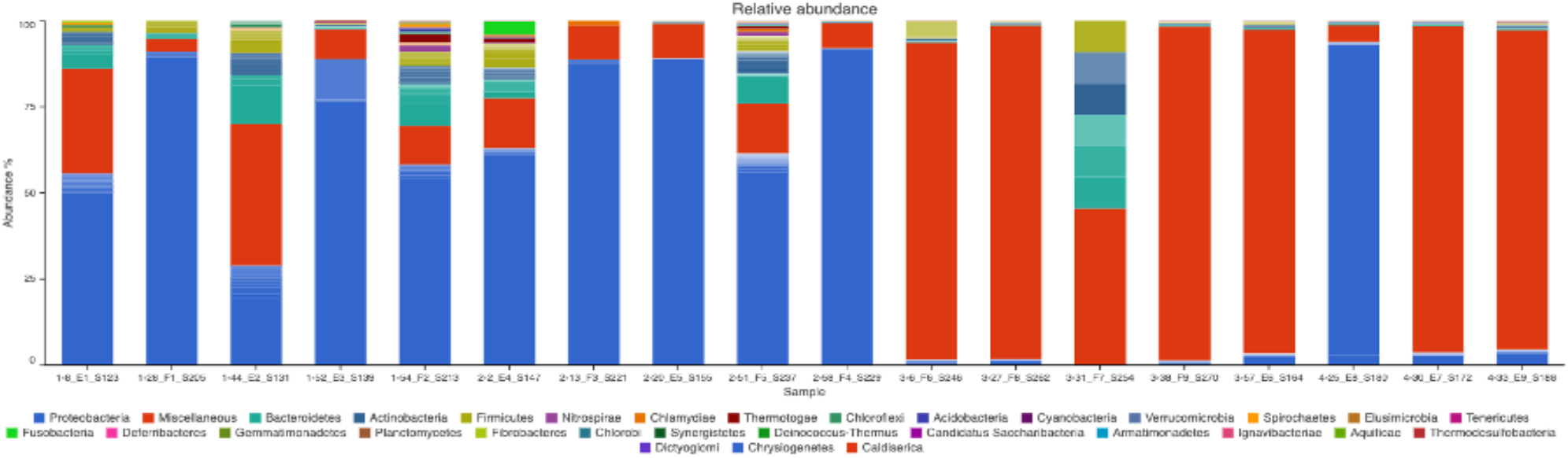
Relative Abundance of OTUs on 7 dpi for 18 samples. The first number before each sample ID identifies the treatment type (1 = No infection, 2 = Infection only, 3 = Infection ZA inject, and 4 = Infection AB inject).

We identify that Fig. 12, illustrates the Relative Abundance of OTU profiles on 7 dpi that follow the approximate total “Mean” profile percentage of the most abundant phylum being *Proteobacteria* (∼84.16%) (Table 7).

**Table 7.**
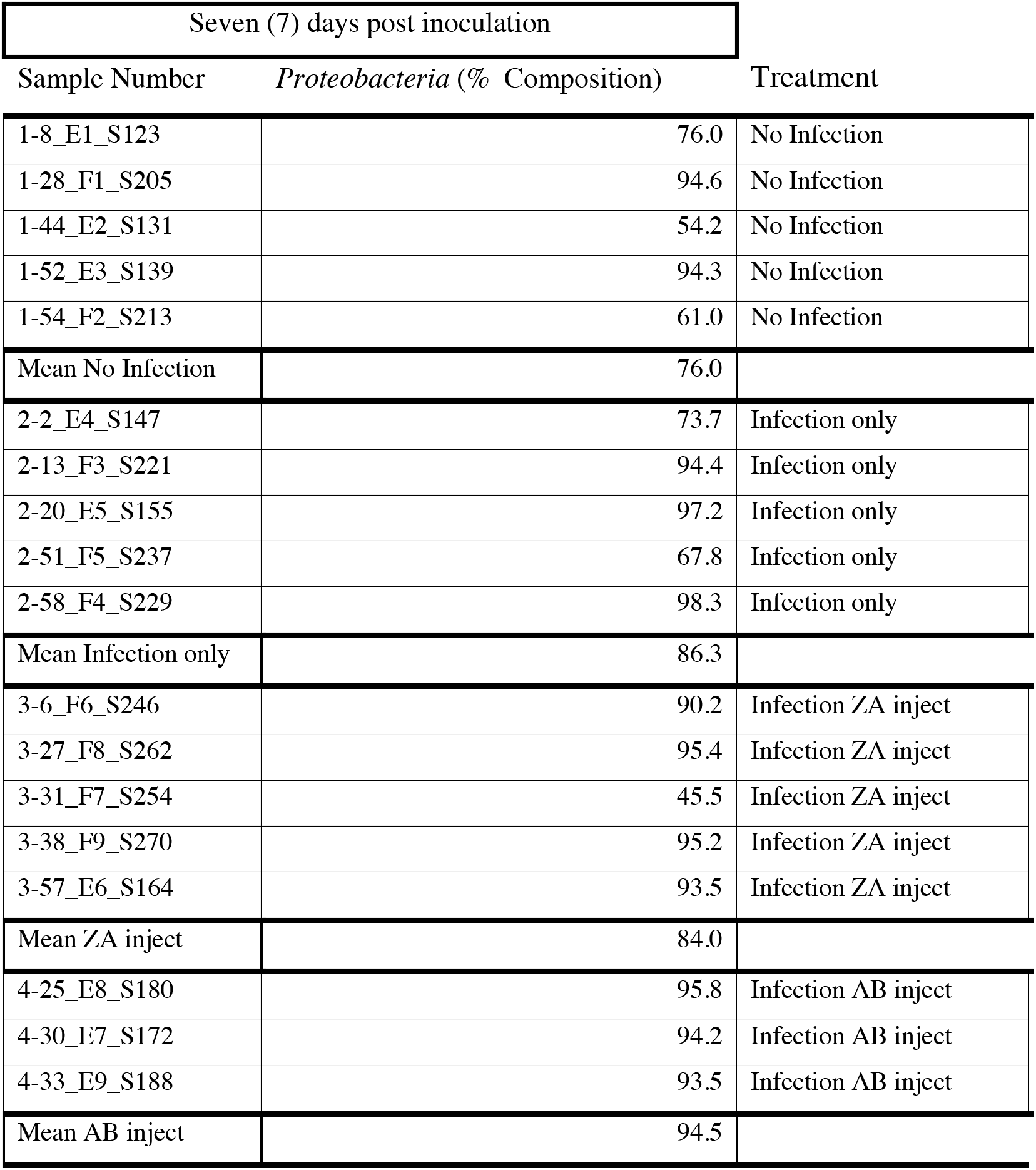
OTU percentage composition. OTU percentage composition of the most prevalent Phylum after 7 days post inoculation (Proteobacteria).

**Table 8.**
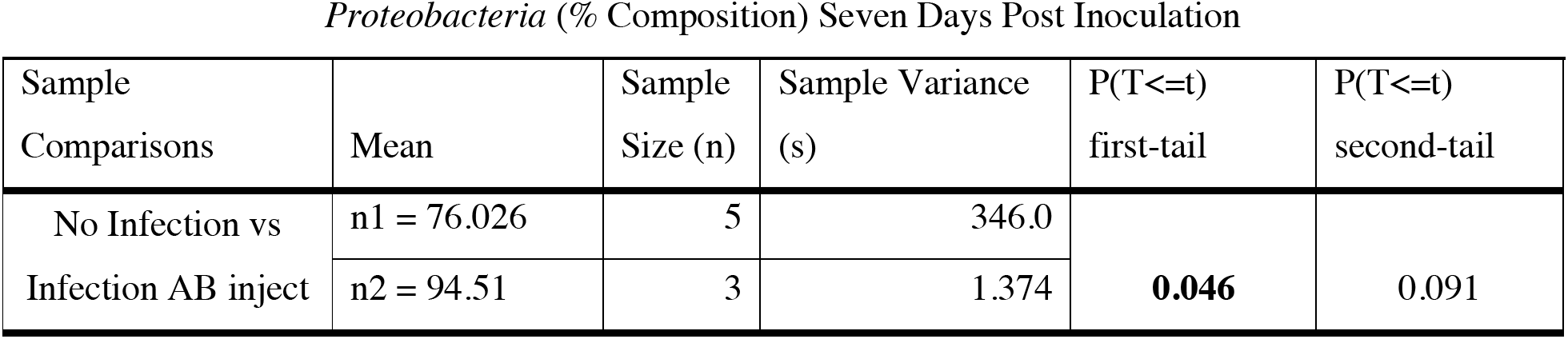
Statistical comparison of Proteobacteria of 7 dpi. Statistical comparison of *Proteobacteria* (% Composition) after seven (7) dpi in host *Capra hircus* wethers.

Gene amplicons for 16S rRNA Relative Abundance of OTU profiles are illustrated for 20 individuals on 21 dpi (Fig. 13). The blood samples are dominated by 36 most abundant OTUs.

**Fig. 13.**
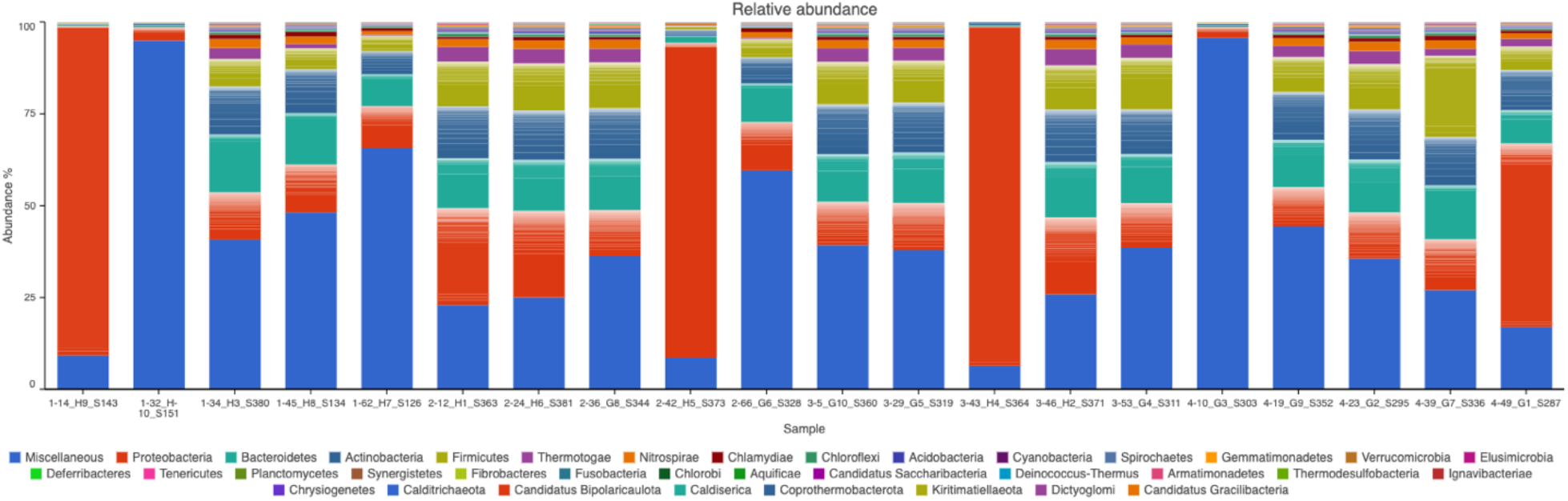
Relative Abundance of OTUs on 21 dpi for 20 samples. The first number before each sample ID identifies the treatment type (1 = No infection, 2 = Infection only, 3 = Infection ZA inject, and 4 = Infection AB inject).

The 21 dpi blood samples accounted for a much more quantified and rich pool of OTUs compared to that of 7 dpi blood samples.

There is clear evidence that greater amounts of difference are present when prolonged exposure to *H. contortus* infection are examined (7 dpi No Infection versus 21 dpi of all other treatments). *Proteobacteria* (Table 10: No Infection 7 dpi vs Infection only 21 dpi; No Infection 7 dpi vs Infection ZA inject 21 dpi; and No Infection 7 dpi vs Infection AB inject 21 dpi) shows the most significant values for relative abundance of OTUs.

**Table 9.**
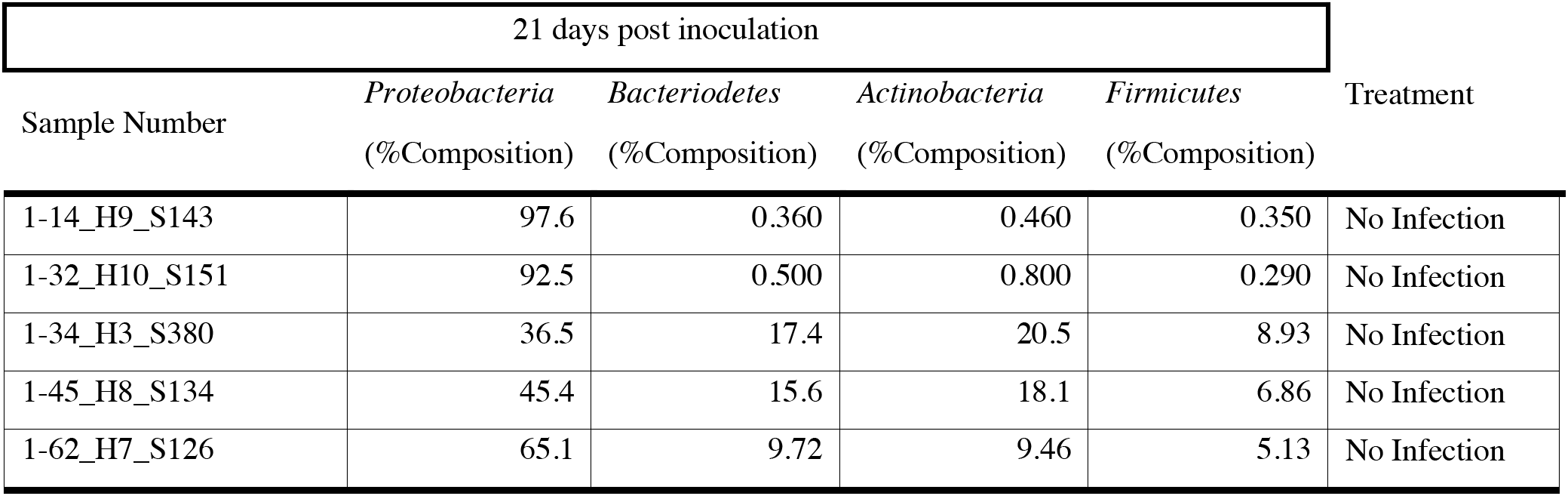
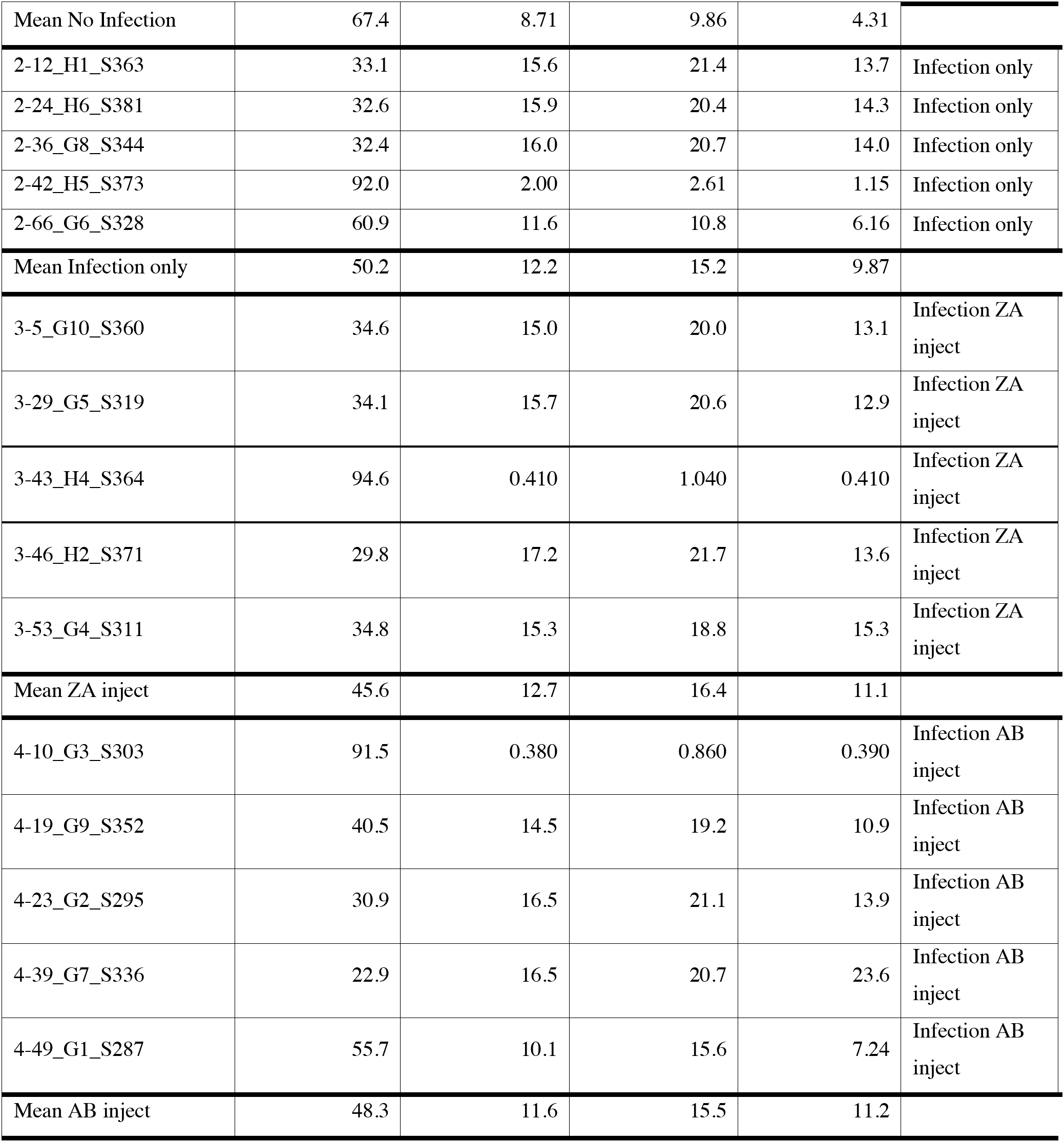
OTU percentage composition of 21 dpi. OTU percentage composition of the most prevalent Phylums after 21 days post inoculation (*Proteobacteria*, *Bacteriodetes*, *Actinobacteria*, and *Firmicutes*).

**Table 10.**
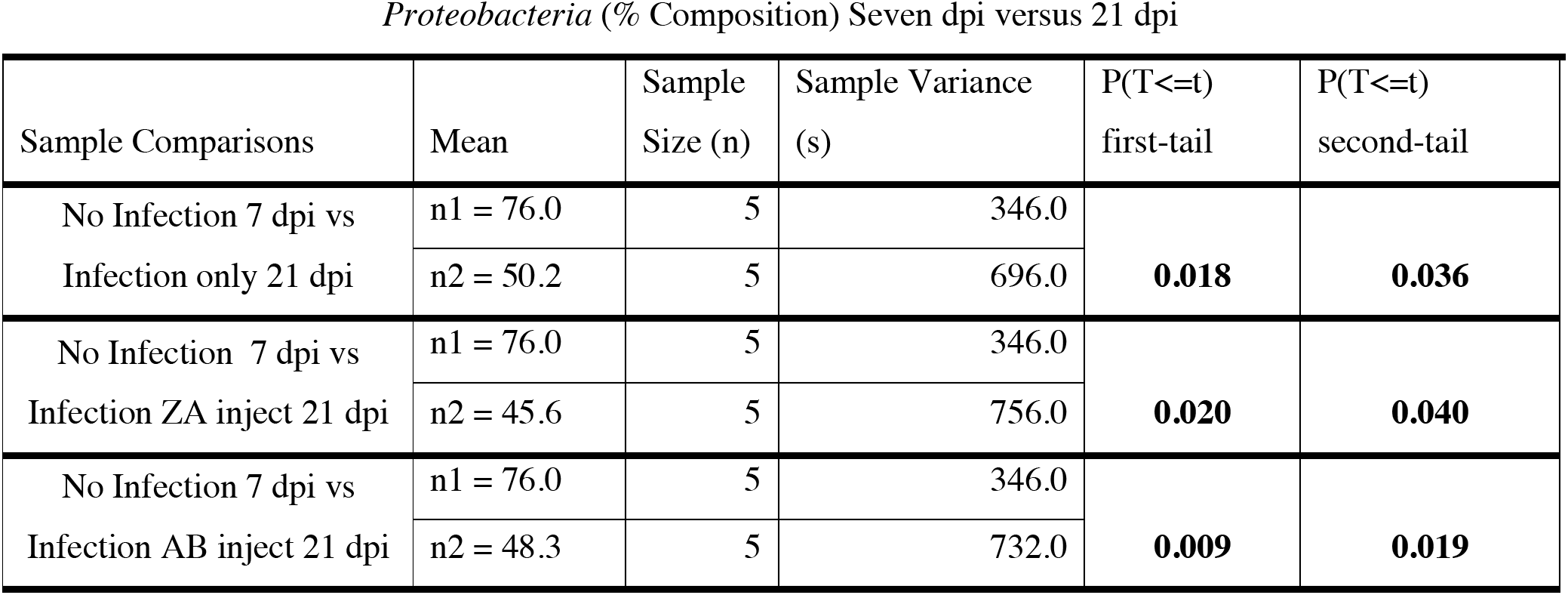
Statistical comparison of Proteobacteria between 7 dpi and 21 dpi. Statistical comparison of *Proteobacteria* (% Composition) after seven (7) dpi versus 21 dpi of host *Capra hircus* wethers.

**Table 11.**
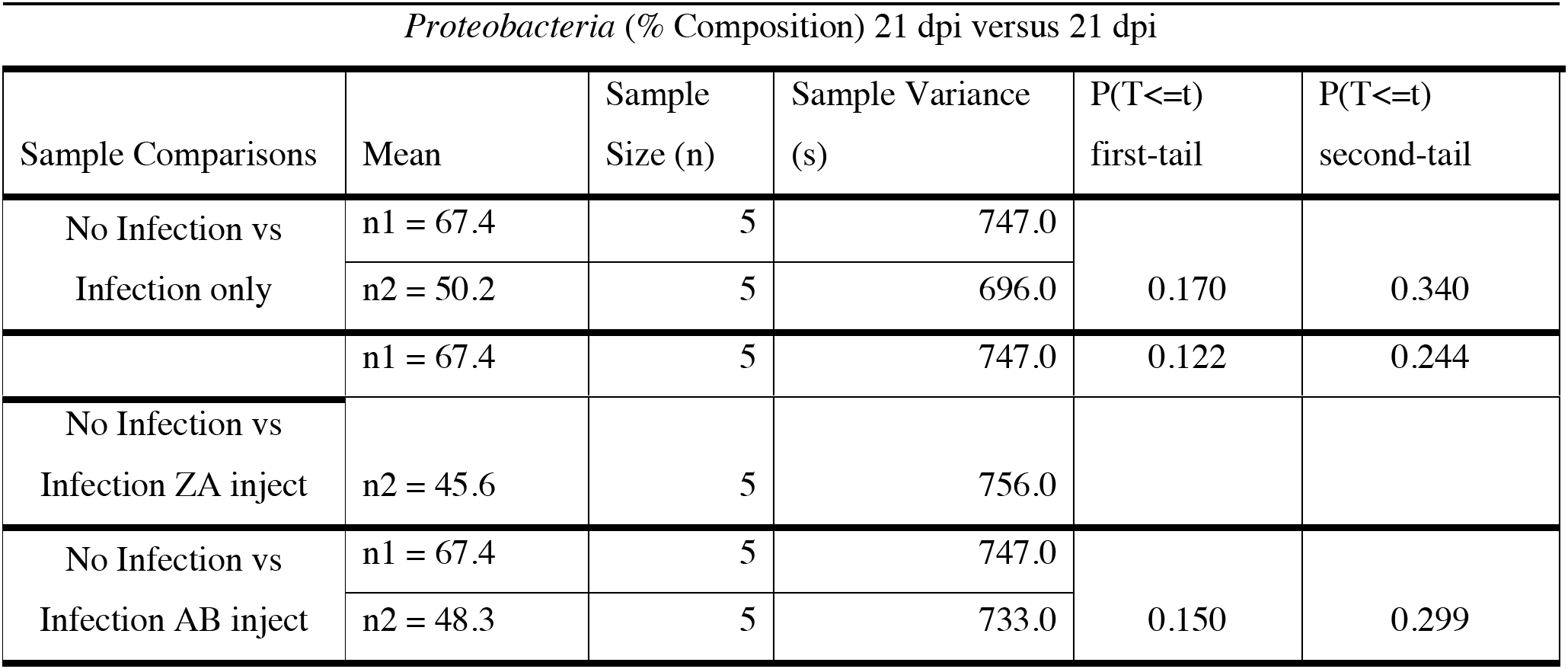
Statistical comparison of *Proteobacteria* of 21 dpi. Statistical comparison of *Proteobacteria* (% Composition) after 21 dpi of host *Capra hircus* wethers.

| Sample Comparisons | Mean | Sample Size (n) | Sample Variance (s) | P(T<=t) first-tail | P(T<=t) second-tail |
| --- | --- | --- | --- | --- | --- |
| No Infection vs Infection only | n1 = 67.4 | 5 | 747.0 | 0.170 | 0.340 |
|  | n2 = 50.2 | 5 | 696.0 |  |  |
|  | n1 = 67.4 | 5 | 747.0 | 0.122 | 0.244 |
| No Infection vs<br>Infection ZA inject | n2 = 45.6 | 5 | 756.0 |  |  |
| No Infection vs<br>Infection AB inject | n1 = 67.4 | 5 | 747.0 | 0.150 | 0.299 |
|  | n2 = 48.3 | 5 | 733.0 |  |  |

**Table 12.**
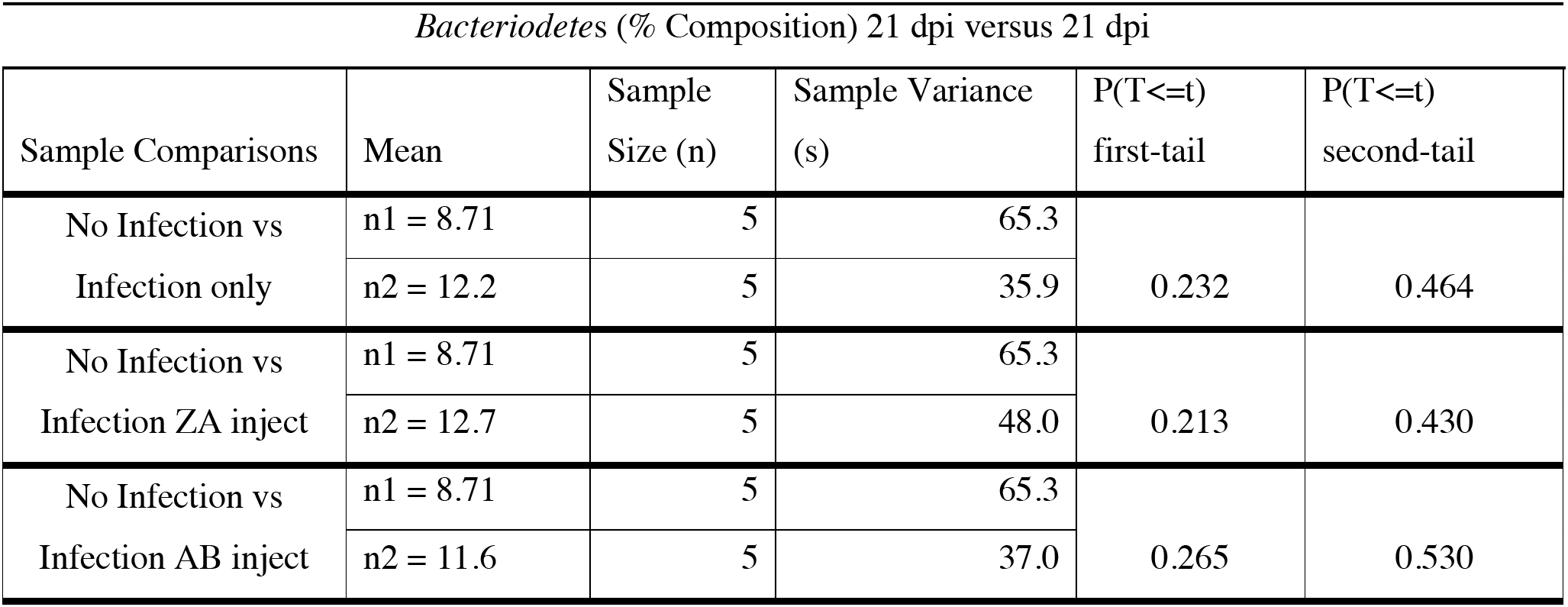
Statistical comparison of Bacteriodetes of 21 dpi. Statistical comparison of *Bacteriodete*s (% Composition) after 21 dpi of host *Capra hircus* wethers

**Table 13.**
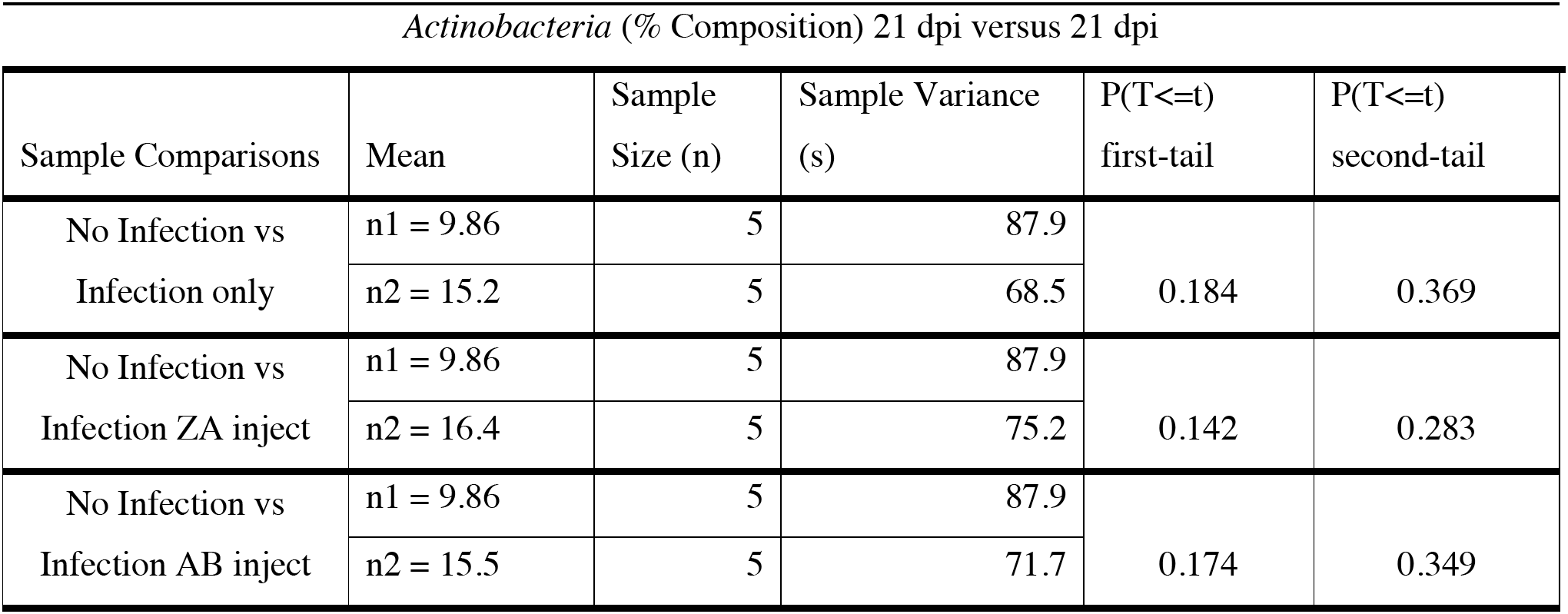
Statistical comparison of *Actinobacteria* of 21 dpi. Statistical comparison of *Actinobacteria* (% Composition) after 21 dpi of host *Capra hircus* wethers.

When a comparison of % Composition were made among 21 dpi treatments, there were P(T<=t) first-tail values that were significant/near significant for *Firmicutes* (Table 14). This implies agreement with the occurrence of inflammation [17] during parasite intrusion.

**Table 14.**
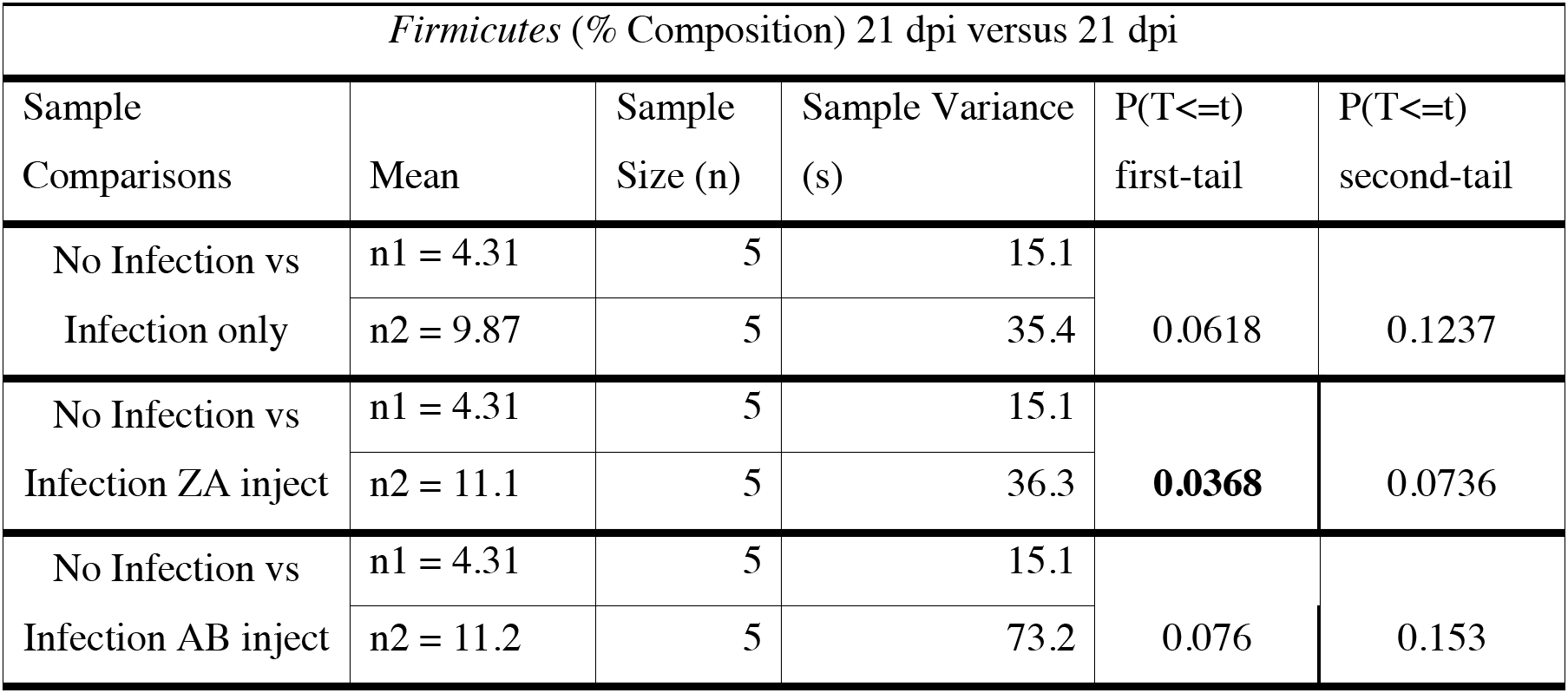
Statistical comparison of Firmicutes of 21 dpi. Statistical comparison of *Firmicutes* (% Composition) after 21 dpi of host *Capra hircus* wethers.

The “Mean” profile percentages of the most abundant phyla shown in Table 9 for “No Infection” were *Proteobacteria* (∼67.4%), followed by *Actinobacteria* (∼9.86%), *Bacteriodetes* (∼8.71%), and *Firmicutes* (∼4.31%) of all OTUs. The “Mean” profile percentages of the most abundant phyla shown in Table 9 for “Infection only” were *Proteobacteria* (∼50.2%), followed by *Actinobacteria* (∼15.2%), *Bacteriodetes* (∼12.2%), and *Firmicutes* (∼9.87%) of all OTUs. The “Mean” profile percentages of the most abundant phyla shown in Table 9 for infection ZA inject were *Proteobacteria* (∼45.6%), followed by *Actinobacteria* (∼16.4%), *Bacteriodetes* (∼12.7%), and *Firmicutes* (∼11.1%) of all OTUs. The “Mean” profile percentages of the most abundant phyla shown in Table 9 for Infection AB inject were *Proteobacteria* (∼48.3%), followed by *Actinobacteria* (∼15.5%), *Bacteriodetes* (∼11.6%), and *Firmicutes* (∼11.2%) of all OTUs.

Table 15 shows the most statistically significant treatment values for *Firmicutes* % Composition/*Bacteriodetes* % Composition (*F*/*B*) ratios. When comparing No Infection versus the other treatments, statistical comparisons revealed that two treatments indicated significant differences (* = p < 0.05): No Infection 21 dpi vs Infection ZA inject 21 dpi and No Infection 21 dpi vs Infection AB inject 21 dpi for the P(T<=t) first-tail.

**Table 15.**
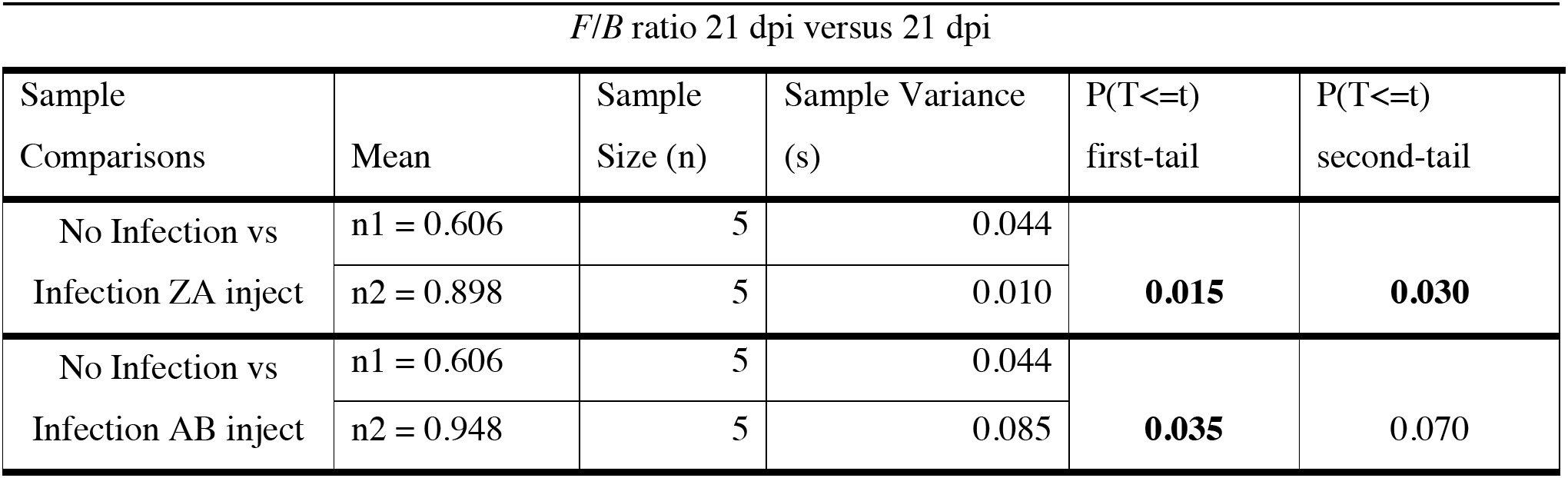
Statistical comparison of *F*/*B* ratios. Statistical comparison of *F*/*B* ratios % Composition after 21 dpi in host *Capra hircus* wethers.

The non-infected controls, after 21 dpi, revealed both less *Bacteriodetes* % Composition and *Firmicutes* % Composition than did all the other treatments. There is a 0.494 ratio for the “Mean” *Bacteriodetes* % Composition to *Firmicutes* % Composition, when non-infected controls are examined. The “Infection only” treatments had a 0.809 ratio for the “Mean” *Bacteriodetes* % Composition to *Firmicutes* % Composition. The treated ZA inject had a ratio of 0.874 and the treated AB inject had a ratio of 0.966 “Mean” *Bacteriodetes* % Composition to *Firmicutes* % Composition. This supports that there are measurable outcomes that can differentiate metabolic healthy wethers from parasitically compromised wethers based on changes in *Bacteriodetes* % Composition to *Firmicutes* % Composition [17].

## Discussion

To characterize the nematode-infected host-transcriptome, host-microbiome, we examined the distribution of 19-7 dpi transcriptome sequencing (RNA-Seq; n = 40) samples and microbial flora from 18-seven (7) dpi and 20-21 dpi samples of four treatment types of 16S rRNA sequencing (MiSeq, n= 80). We focused on whole-blood or buffy coat derived extractions of mbDNA [26,27,28] to discriminate among non-infected, metabolically healthy controls and infected types. We performed microbial assignments using Kraken [29] and transcriptome analysis using STAR [30], completing differential analysis using GSA [31]. Our data identified 7627 genes that were expressed and shows differences in association across treatment types on 7 dpi and 21 dpi in specific microbiota profiles.

The evidence determined in this study shows that blood-based microbial DNA (mbDNA) can be used to discriminate between a non-infected, metabolically healthy or a *Haemonchus contortus-*attacked *Capra hircus wethers* [32]. The evidence also shows that the microbiome changes in abundance and diversity over age (dpi), even within the interval of a few days [33,34,35,36,37], further supporting that microbial flora abundance and diversity is associated with metabolic health or the indication of non-infection [44].

Factors, including antibiotics, can change the composition of microbiota [38,39,40,41] often destroying the composition of beneficial microbes along with pathological ones. Thus, causing dysbiosis or the development of unwanted microbes [41,42,43, 44]. Thus, the residual infection condition is still evident despite antibiotic or other type of treatment, indicated by inflammatory results or an increase in ratio of inflammation causing microbial flora. We identified evidence that the composition of inflammatory microbiota increases with zoledronic acid or anti-γσ T cells treatment. With a greater increase being exhibited by anti-γσ T cells treatment.

Unlike the results described by a previous study where it was surmised that infection by *H. contortus* did not affect caprine microbial diversity [8], we identified that in goat blood samples there were likely significant differences based especially on treatment types. There is also likely significance in evidence that microbial abundance and diversity differences are dependent on age (days post inoculation).

Our results imply that there are differences in gene expression as subjects become older or are exposed to the environment longer. Attributes of the different treatment types show that genes are expressed in response to the *H. contortus* presence. All pointing towards the implication of *H. contortus* to effectively change and disrupt the internal habitat of the host.

The host health may be determined based on *Firmicutes*/*Bacteroidetes (F*/*B)* ratios. The *F*/*B* Ratio is estimated by utilizing the lowest and highest values of the reference range (uninfected host) for individual organisms [44]. A high *F*/*B* ratio may be related to increased caloric extraction from food, fat deposition and lipogenesis, impaired insulin sensitivity, and increased inflammation. Low *Firmicutes*/*Bacteroidetes (F*/*B)* ratio, is an indicator of dysbiosis indicated by a decreased diversity of the microbiome compared to healthy cohorts. *F*/*B* is considered as “low” when the value falls below a threshold [44]. Therefore, “metabolic health” is determined by the actions taken by the parasite. If the parasite yields an *F*/*B* ratio that is depressed compared to the reference range of harmful intrusion, it may be postulated that the host does not have adequate means of defending itself, thus the parasite load is depleting host resources. Alternatively, if the parasite yields an *F*/*B* ratio that is elevated compared to the reference range of harmful intrusion, it indicates that the host is being hyperactive or inflamed as a response to the parasite burden.

## Conclusions

The metabolic systems affected warrant further investigation to identify specific pathways where significant changes have resulted based on being exposed to infection. Furthermore, the development of computational algorithms for correlation of microbial abundance and diversity are warranted. The authors conjecture that blood samples are shown here to be a possible means to indicate *H. contortus* infection based on detection of microbial flora abundance and diversity as well as in gene expression profiles. Correlations can be drawn on statistical levels of microbial flora for this specific type of inflammation. The specificity in the range of microbial flora, dependent on the age of the *wethers*, can indicate the occurrence of depleting resources or inflammation due to *H. contortus* infection. In other words, this implies *H. contortus* does effectively change and disrupt the internal habitat health of the host and the effects are measurable. The development of a standard laboratory diagnostic procedure using blood microbiota to detect gastrointestinal infection with *H. contortus* is the ideal course of action.

## Supporting information

Table 1. Experimental Set-up

Table 2. Detailed Experimental Set-up

Table 3. Alpha diversity report for 7 dpi Shannon and Simpson index

Table 4. Significant values of 7 dpi Shannon and Simpson indices

Table 5. Alpha diversity report for 21 dpi Shannon and Simpson in

Table 6. Statistically different comparison results of 7 dpi versus 21 dpi

Table 7. OTU percentage composition

Table 8. Statistical comparison of Proteobacteria of 7 dpi

Table 9. OTU percentage composition of 21 dpi

Table 10. Statistical comparison of Proteobacteria between 7 dpi and 21 dpi

Table 11. Statistical comparison of Proteobacteria of 21 dpi

Table 12. Statistical comparison of Bacteriodetes of 21 dpi

Table 13. Statistical comparison of Actinobacteria of 21 dpi

Table 14. Statistical comparison of Firmicutes of 21 dpi

Table 15. Statistical comparison of F/B ratios

## Availability of data and materials

The metagenomic data have been deposited with links to BioProject accession number PRJNA612987 in the NCBI BioProject database (https://www.ncbi.nlm.nih.gov/bioproject/).

The gene expression data discussed in this publication have been deposited in NCBI’s Gene Expression Omnibus [45] and are accessible through GEO Series accession number GSE169607 (https://www.ncbi.nlm.nih.gov/geo/query/acc.cgi?acc=GSE169607.

## Acknowledgements

The author is thankful to coinvestigators Zaisen Wang, Qunhui Yang, and Jessica Quijada-Pinango and the Director of the American Institute for Goat research (AIGR: School of Agriculture and Applied Sciences, Langston University, Langston, Oklahoma, USA), Mrs. Kelly Kyle and Mrs. Xiaowen Wang (Partek Incorporated). The Relationship Between the Microbiome and Internal Parasitism in Goats is supported by the Evans Allen Program Grant at the Agricultural Research and Extension Center at Langston University.

