## Supplementary material for "Functional genome and microbiome in blood of goats affected by the gastrointestinal pathogen *Haemonchus contortus*": Table 1. Experimental Set-up

**Table 1. Experimental Set-up.** Treatment Group (1-4) for infection L3 *H. contortus* (+) or non-infection *H. contortus* (-), with (+) or without (-) treatment type (Zoledronic acid injection or γδ T depletion).

| Group | L3 *H. contortus* infection | Zoledronic acid | γδ T depletion |
|---|---|---|---|
| 1 | - | - | - |
| 2 | + | - | - |
| 3 | + | + | - |
| 4 | + | - | + |
