## Supplementary material for "Functional genome and microbiome in blood of goats affected by the gastrointestinal pathogen *Haemonchus contortus*": Table 2. Detailed Experimental Set-up

**Table 2. Detailed Experimental Set-up.** The Group number, Treatment type, identifying Tag, days post inoculation (dpi), Age in days, and Body Weight (BW) in kg.

| Group | Treatment | Tag/Sample ID | dpi | Age, days | BW, kg |
|---|---|---|---|---|---|
| 1 | No Infection | 8_E1_S123 | 7 | 118 | 21.05 |
| 1 | No Infection | 28_F1_S205 | 7 | 116 | 17.9 |
| 1 | No Infection | 44_E2_S131 | 7 | 115 | 18.95 |
| 1 | No Infection | 52_E3_S139 | 7 | 113 | 17.75 |
| 1 | No Infection | 54_F2_S213 | 7 | 114 | 22.8 |
| 2 | Infection only | 13_F3_S221 | 7 | 118 | 22.9 |
| 2 | Infection only | 2_E4_S147 | 7 | 120 | 19.05 |
| 2 | Infection only | 20_E5_S155 | 7 | 117 | 16.8 |
| 2 | Infection only | 58_F4_S229 | 7 | 100 | 17.75 |
| 2 | Infection only | 51_F5_S237 | 7 | 113 | 16.65 |
| 3 | Infection ZA inject | 6_F6_S246 | 7 | 119 | 17.65 |
| 3 | Infection ZA inject | 31_F7_S254 | 7 | 116 | 17.7 |
| 3 | Infection ZA inject | 27_F8_S262 | 7 | 116 | 20.9 |
| 3 | Infection ZA inject | 38_F9_S270 | 7 | 115 | 20.35 |
| 3 | Infection ZA inject | 57_E6_S164 | 7 | 96 | 19.7 |
| 4 | Infection AB inject | 17_F10_S18 | 7 | 117 | 16.8 |
| 4 | Infection AB inject | 25_E8_S180 | 7 | 116 | 19.95 |
| 4 | Infection AB inject | 33_E9_S188 | 7 | 116 | 19.8 |
| 4 | Infection AB inject | 30_E7_S172 | 7 | 116 | 19.7 |
| 4 | Infection AB inject | 50_E10_S9 | 7 | 114 | 21.65 |
