## Supplementary material for "Functional genome and microbiome in blood of goats affected by the gastrointestinal pathogen *Haemonchus contortus*": Table 3. Alpha diversity report for 7 dpi Shannon and Simpson index

| Sample name | Shannon index | Simpson index | dpi | treatment |
|---|---|---|---|---|
| 1-8_E1_S123 | 1.05 | 0.360 | 7 | No Infection |
| 1-28_F1_S205 | 0.190 | 0.0747 | 7 | No Infection |
| 1-44_E2_S131 | 2.05 | 0.772 | 7 | No Infection |
| 1-52_E3_S139 | 0.492 | 0.254 | 7 | No Infection |
| 1-54_F2_S213 | 1.49 | 0.487 | 7 | No Infection |
| Mean No Infection | 1.06 | 0.3894 | | |
| 2-13_F3_S221 | 0.1607 | 0.061 | 7 | Infection only |
| 2-2_E4_S147 | 1.40 | 0.401 | 7 | Infection only |
| 2-20_E5_S155 | 0.0866 | 0.0198 | 7 | Infection only |
| 2-58_F4_S229 | 0.0535 | 0.0133 | 7 | Infection only |
| 2-51_F5_S237 | 1.828 | 0.486 | 7 | Infection only |
| Mean Infection only | 0.707 | 0.196 | | |
| 3-6_F6_S246 | 1.58 | 0.591 | 7 | Infection ZA inject |
| 3-31_F7_S254 | 1.61 | 0.800 | 7 | Infection ZA inject |
| 3-27_F8_S262 | 2.86 | 0.823 | 7 | Infection ZA inject |
| 3-38_F9_S270 | 3.52 | 0.940 | 7 | Infection ZA inject |
| 3-57_E6_S164 | 2.77 | 0.757 | 7 | Infection ZA inject |
| Mean Infection ZA inject | 2.47 | 0.782 | | |
| 4-25_E8_S180 | 0.317 | 0.099 | 7 | Infection AB inject |
| 4-33_E9_S188 | 2.64 | 0.733 | 7 | Infection AB inject |
| 4-30_E7_S172 | 2.35 | 0.659 | 7 | Infection AB inject |
| Mean Infection AB inject | 1.770 | 0.497 | | |
