## Supplementary material for "Functional genome and microbiome in blood of goats affected by the gastrointestinal pathogen *Haemonchus contortus*": Table 4. Significant values of 7 dpi Shannon and Simpson indices

**Table 4. Significant values of 7 dpi Shannon and Simpson indices.** Statistical comparison of a) Shannon and b) Simpson distribution indices for Alpha diversity reports of 7 dpi richness and diversity of microbial flora in host *Capra hircus* wethers.

a) Shannon index

| Sample Comparisons | Mean | Sample Size (n) | Sample Variance (s) | P(T<=t) first-tail | P(T<=t) second-tail |
|---|---|---|---|---|---|
| No Infection vs Infection ZA inject | n1 = 1.06 | 5 | 0.562 | **0.012** | **0.023** |
| | n2 = 2.47 | 5 | 0.718 | | |

b) Simpson index

| Sample Comparisons | Mean | Sample Size (n) | Sample Variance (s) | P(T<=t) first-tail | P(T<=t) second-tail |
|---|---|---|---|---|---|
| No Infection vs Infection ZA inject | n1 = 0.389 | 5 | 0.0685 | **0.012** | **0.023** |
| | n2 = 0.782 | 5 | 0.0160 | | |
