## Supplementary material for "Functional genome and microbiome in blood of goats affected by the gastrointestinal pathogen *Haemonchus contortus*": Table 5. Alpha diversity report for 21 dpi Shannon and Simpson in

**Table 5. Alpha diversity report for 21 dpi Shannon and Simpson index.**

| Sample name | Shannon index | Simpson index | dpi | treatment |
|---|---|---|---|---|
| 1-14_H9_S143 | 0.240 | 0.0643 | 21 | No Infection |
| 1-32_H10_S151 | 2.64 | 0.729 | 21 | No Infection |
| 1-45_H8_S134 | 3.83 | 0.901 | 21 | No Infection |
| 1-34_H3_S380 | 4.20 | 0.911 | 21 | No Infection |
| 1-62_H7_S126 | 3.63 | 0.892 | 21 | No Infection |
| Mean No Infection | 2.91 | 0.699 | | |
| 2-12_H1_S363 | 3.97 | 0.917 | 21 | Infection only |
| 2-24_H6_S381 | 4.21 | 0.925 | 21 | Infection only |
| 2-36_G8_S344 | 4.71 | 0.949 | 21 | Infection only |
| 2-42_H5_S373 | 0.461 | 0.113 | 21 | Infection only |
| 2-66_G6_S328 | 3.52 | 0.886 | 21 | Infection only |
| Mean Infection only | 3.37 | 0.758 | | |
| 3-5_G10_S360 | 4.68 | 0.952 | 21 | Infection ZA inject |
| 3-29_G5_S319 | 4.72 | 0.950 | 21 | Infection ZA inject |
| 3-43_H4_S364 | 0.145 | 0.0438 | 21 | Infection ZA inject |
| 3-53_G4_S311 | 4.38 | 0.929 | 21 | Infection ZA inject |
| 3-46_H2_S371 | 4.31 | 0.934 | 21 | Infection ZA inject |
| Mean Infection ZA inject | 3.65 | 0.762 | | |
| 4-10_G3_S303 | 2.76 | 0.789 | 21 | infection AB inject |
| 4-19_G9_S352 | 4.66 | 0.950 | 21 | infection AB inject |
| 4-23_G2_S295 | 4.83 | 0.958 | 21 | infection AB inject |
| 4-39_G7_S336 | 3.62 | 0.886 | 21 | infection AB inject |
| 4-49_G1_S287 | 2.51 | 0.644 | 21 | infection AB inject |
| Mean Infection AB inject | 3.68 | 0.845 | | |
