## Supplementary material for "Functional genome and microbiome in blood of goats affected by the gastrointestinal pathogen *Haemonchus contortus*": Table 6. Statistically different comparison results of 7 dpi versus 21 dpi

**Table 6. Statistically different comparison results of 7 dpi versus 21 dpi.** Statistically different comparison results of 7 dpi versus 21 dpi a) Shannon and b) Simpson distribution indices for Alpha diversity reports of abundance and diversity for microbial flora in host *Capra hircus* wethers.

a) Shannon index 7 dpi vs 21 dpi t-Test: Two-Sample Assuming Unequal Variances

| Sample Comparisons | Mean | Sample Size (n) | Sample Variance (s) | P(T<=t) one-tail | P(T<=t) two-tail |
|---|---|---|---|---|---|
| No Infection 7 dpi | n1 = 1.05 | 5 | 0.562 | | |
| vs No Infection 21 dpi | n2 =2.91 | 5 | 2.56 | **0.029** | 0.057 |
| No Infection 7 dpi | n1 =1.05 | 5 | 0.562 | | |
| vs Infection only 21 dpi | n2 = 3.37 | 5 | 2.83 | **0.015** | **0.031** |
| No Infection 7 dpi | n1 = 1.05 | 5 | 0.562 | | |
| vs Infection ZA inject 21 dpi | n2 = 3.65 | 5 | 3.86 | **0.020** | **0.040** |
| No Infection 7 dpi | n1 =1.05 | 5 | 0.562 | | |
| vs Infection AB inject 21 dpi | n2 = 3.67 | 5 | 1.12 | **0.001** | **0.003** |
| Infection only 7 dpi | n1 = 0.707 | 5 | 0.714 | | |
| vs No Infection 21 dpi | n2 = 2.91 | 5 | 2.56 | **0.017** | **0.035** |
| Infection only 7 dpi | n1 = 0.707 | 5 | 0.714 | | |
| vs Infection only 21 dpi | n2 = 3.37 | 5 | 2.83 | **0.010** | **0.020** |
| Infection only 7 dpi | n1 = 0.707 | 5 | 0.714 | | |
| vs | n2 = 3.65 | 5 | 3.86 | **0.014** | **0.028** |

| | | | | | |
|---|---|---|---|---|---|
| Infection ZA inject 21 dpi | | | | | |
| Infection only 7dpi vs Infection AB inject 21 dpi | n1 = 0.707 | 5 | 0.714 | | |
| | n2 = 3.67 | 5 | 1.12 | **0.001** | **0.001** |
| Infection ZA inject 7dpi vs Infection AB inject 21 dpi | n1 = 2.47 | 5 | 0.718 | | |
| | n2 = 3.67 | 5 | 1.12 | **0.041** | 0.082 |
| Infection AB inject 7dpi vs Infection AB inject 21 dpi | n1 = 1.77 | 3 | 1.60 | | |
| | n2 = 3.67 | 5 | 1.12 | **0.047** | 0.094 |

Simpson index 7dpi vs 21 dpi t-Test: Two-Sample Assuming Unequal Variances

| Sample Comparisons | Mean | Sample Size (n) | Sample Variance (s) | P(T<=t) one-tail | P(T<=t) two-tail |
|---|---|---|---|---|---|
| No Infection 7 dpi vs Infection AB inject 21 dpi | n1 = 0.390 | 5 | 0.0685 | | |
| | n2 = 0.845 | 5 | 0.0172 | **0.007** | **0.013** |
| Infection only 7 dpi vs No Infection 21 dpi | n1 = 0.196 | 5 | 0.0522 | | |
| | n2 = 0.699 | 5 | 0.132 | **0.017** | **0.034** |
| Infection only 7 dpi vs Infection only 21 dpi | n1 = 0.196 | 5 | 0.0522 | | |
| | n2 = 0.758 | 5 | 0.130 | **0.011** | **0.022** |
| Infection only 7 dpi vs Infection ZA inject 21 dpi | n1 = 0.196 | 5 | 0.0522 | | |
| | n2 = 0.762 | 5 | 0.161 | **0.017** | **0.034** |

| | | | | | |
|---|---|---|---|---|---|
| Infection only 7dpi vs Infection AB inject 21 dpi | n1 = 0.196 | 5 | 0.0522 | **0.001** | **0.002** |
