## Supplementary material for "Functional genome and microbiome in blood of goats affected by the gastrointestinal pathogen *Haemonchus contortus*": Table 7. OTU percentage composition

**Table 7. OTU percentage composition.** OTU percentage composition of the most prevalent Phylum after 7 days post inoculation (Proteobacteria).

| Seven (7) days post inoculation | | |
|---|---|---|
| Sample Number | *Proteobacteria* (% Composition) | Treatment |
| 1-8_E1_S123 | 76.0 | No Infection |
| 1-28_F1_S205 | 94.6 | No Infection |
| 1-44_E2_S131 | 54.2 | No Infection |
| 1-52_E3_S139 | 94.3 | No Infection |
| 1-54_F2_S213 | 61.0 | No Infection |
| Mean No Infection | 76.0 | |
| 2-2_E4_S147 | 73.7 | Infection only |
| 2-13_F3_S221 | 94.4 | Infection only |
| 2-20_E5_S155 | 97.2 | Infection only |
| 2-51_F5_S237 | 67.8 | Infection only |
| 2-58_F4_S229 | 98.3 | Infection only |
| Mean Infection only | 86.3 | |
| 3-6_F6_S246 | 90.2 | Infection ZA inject |
| 3-27_F8_S262 | 95.4 | Infection ZA inject |
| 3-31_F7_S254 | 45.5 | Infection ZA inject |
| 3-38_F9_S270 | 95.2 | Infection ZA inject |
| 3-57_E6_S164 | 93.5 | Infection ZA inject |
| Mean ZA inject | 84.0 | |
| 4-25_E8_S180 | 95.8 | Infection AB inject |
| 4-30_E7_S172 | 94.2 | Infection AB inject |
| 4-33_E9_S188 | 93.5 | Infection AB inject |
| Mean AB inject | 94.5 | |
