## Supplementary material for "Functional genome and microbiome in blood of goats affected by the gastrointestinal pathogen *Haemonchus contortus*": Table 8. Statistical comparison of Proteobacteria of 7 dpi

**Table 8. Statistical comparison of *Proteobacteria* of 7 dpi.** Statistical comparison of *Proteobacteria* (% Composition) after seven (7) dpi in host *Capra hircus* wethers.

*Proteobacteria* (% Composition) Seven Days Post Inoculation

| Sample Comparisons | Mean | Sample Size (n) | Sample Variance (s) | P(T<=t) first-tail | P(T<=t) second-tail |
|---|---|---|---|---|---|
| No Infection vs Infection AB inject | n1 = 76.026 | 5 | 346.0 | **0.046** | 0.091 |
| | n2 = 94.51 | 3 | 1.374 | | |
