## Supplementary material for "Functional genome and microbiome in blood of goats affected by the gastrointestinal pathogen *Haemonchus contortus*": Table 9. OTU percentage composition of 21 dpi

**Table 9. OTU percentage composition of 21 dpi.** OTU percentage composition of the most prevalent Phylums after 21 days post inoculation (*Proteobacteria*, *Bacteriodetes*, *Actinobacteria*, and *Firmicutes*).

| 21 days post inoculation | | | | | |
|---|---|---|---|---|---|
| Sample Number | *Proteobacteria* (%Composition) | *Bacteriodetes* (%Composition) | *Actinobacteria* (%Composition) | *Firmicutes* (%Composition) | Treatment |
| 1-14_H9_S143 | 97.6 | 0.360 | 0.460 | 0.350 | No Infection |
| 1-32_H10_S151 | 92.5 | 0.500 | 0.800 | 0.290 | No Infection |
| 1-34_H3_S380 | 36.5 | 17.4 | 20.5 | 8.93 | No Infection |
| 1-45_H8_S134 | 45.4 | 15.6 | 18.1 | 6.86 | No Infection |
| 1-62_H7_S126 | 65.1 | 9.72 | 9.46 | 5.13 | No Infection |
| Mean No Infection | 67.4 | 8.71 | 9.86 | 4.31 | |
| 2-12_H1_S363 | 33.1 | 15.6 | 21.4 | 13.7 | Infection only |
| 2-24_H6_S381 | 32.6 | 15.9 | 20.4 | 14.3 | Infection only |
| 2-36_G8_S344 | 32.4 | 16.0 | 20.7 | 14.0 | Infection only |
| 2-42_H5_S373 | 92.0 | 2.00 | 2.61 | 1.15 | Infection only |
| 2-66_G6_S328 | 60.9 | 11.6 | 10.8 | 6.16 | Infection only |
| Mean Infection only | 50.2 | 12.2 | 15.2 | 9.87 | |
| 3-5_G10_S360 | 34.6 | 15.0 | 20.0 | 13.1 | Infection ZA inject |
| 3-29_G5_S319 | 34.1 | 15.7 | 20.6 | 12.9 | Infection ZA inject |
| 3-43_H4_S364 | 94.6 | 0.410 | 1.040 | 0.410 | Infection ZA inject |
| 3-46_H2_S371 | 29.8 | 17.2 | 21.7 | 13.6 | Infection ZA inject |
| 3-53_G4_S311 | 34.8 | 15.3 | 18.8 | 15.3 | Infection ZA inject |
| Mean ZA inject | 45.6 | 12.7 | 16.4 | 11.1 | |
| 4-10_G3_S303 | 91.5 | 0.380 | 0.860 | 0.390 | Infection AB inject |
| 4-19_G9_S352 | 40.5 | 14.5 | 19.2 | 10.9 | Infection AB inject |

| | | | | | |
|---|---|---|---|---|---|
| 4-23_G2_S295 | 30.9 | 16.5 | 21.1 | 13.9 | Infection AB inject |
| 4-39_G7_S336 | 22.9 | 16.5 | 20.7 | 23.6 | Infection AB inject |
| 4-49_G1_S287 | 55.7 | 10.1 | 15.6 | 7.24 | Infection AB inject |
| Mean AB inject | 48.3 | 11.6 | 15.5 | 11.2 | |
