## Supplementary material for "Functional genome and microbiome in blood of goats affected by the gastrointestinal pathogen *Haemonchus contortus*": Table 10. Statistical comparison of Proteobacteria between 7 dpi and 21 dpi

**Table 10. Statistical comparison of *Proteobacteria* between 7 dpi and 21 dpi.** Statistical comparison of *Proteobacteria* (% Composition) after seven (7) dpi versus 21 dpi of host *Capra hircus* wethers.

*Proteobacteria* (% Composition) Seven dpi versus 21 dpi

| Sample Comparisons | Mean | Sample Size (n) | Sample Variance (s) | P(T<=t) first-tail | P(T<=t) second-tail |
|---|---|---|---|---|---|
| No Infection 7 dpi vs Infection only 21 dpi | n1 = 76.0 | 5 | 346.0 | **0.018** | **0.036** |
| | n2 = 50.2 | 5 | 696.0 | | |
| No Infection 7 dpi vs Infection ZA inject 21 dpi | n1 = 76.0 | 5 | 346.0 | **0.020** | **0.040** |
| | n2 = 45.6 | 5 | 756.0 | | |
| No Infection 7 dpi vs Infection AB inject 21 dpi | n1 = 76.0 | 5 | 346.0 | **0.009** | **0.019** |
| | n2 = 48.3 | 5 | 732.0 | | |
