## Supplementary material for "Functional genome and microbiome in blood of goats affected by the gastrointestinal pathogen *Haemonchus contortus*": Table 11. Statistical comparison of Proteobacteria of 21 dpi

**Table 11. Statistical comparison of *Proteobacteria* of 21 dpi.** Statistical comparison of *Proteobacteria* (% Composition) after 21 dpi of host *Capra hircus* wethers.

| *Proteobacteria* (% Composition) 21 dpi versus 21 dpi | | | | | |
|---|---|---|---|---|---|
| Sample Comparisons | Mean | Sample Size (n) | Sample Variance (s) | P(T<=t) first-tail | P(T<=t) second-tail |
| No Infection vs Infection only | n1 = 67.4 | 5 | 747.0 | 0.170 | 0.340 |
| | n2 = 50.2 | 5 | 696.0 | | |
| No Infection vs Infection ZA inject | n1 = 67.4 | 5 | 747.0 | 0.122 | 0.244 |
| | n2 = 45.6 | 5 | 756.0 | | |
| No Infection vs Infection AB inject | n1 = 67.4 | 5 | 747.0 | 0.150 | 0.299 |
| | n2 = 48.3 | 5 | 733.0 | | |
