## Supplementary material for "Functional genome and microbiome in blood of goats affected by the gastrointestinal pathogen *Haemonchus contortus*": Table 12. Statistical comparison of Bacteriodetes of 21 dpi

**Table 12. Statistical comparison of *Bacteriodetes* of 21 dpi.** Statistical comparison of *Bacteriodetes* (% Composition) after 21 dpi of host *Capra hircus* wethers

| *Bacteriodetes* (% Composition) 21 dpi versus 21 dpi | | | | | |
|---|---|---|---|---|---|
| Sample Comparisons | Mean | Sample Size (n) | Sample Variance (s) | P(T<=t) first-tail | P(T<=t) second-tail |
| No Infection vs Infection only | n1 = 8.71 | 5 | 65.3 | 0.232 | 0.464 |
| | n2 = 12.2 | 5 | 35.9 | | |
| No Infection vs Infection ZA inject | n1 = 8.71 | 5 | 65.3 | 0.213 | 0.430 |
| | n2 = 12.7 | 5 | 48.0 | | |
| No Infection vs Infection AB inject | n1 = 8.71 | 5 | 65.3 | 0.265 | 0.530 |
| | n2 = 11.6 | 5 | 37.0 | | |
