## Supplementary material for "Functional genome and microbiome in blood of goats affected by the gastrointestinal pathogen *Haemonchus contortus*": Table 13. Statistical comparison of Actinobacteria of 21 dpi

**Table 13. Statistical comparison of *Actinobacteria* of 21 dpi.** Statistical comparison of *Actinobacteria* (% Composition) after 21 dpi of host *Capra hircus* wethers.

| *Actinobacteria* (% Composition) 21 dpi versus 21 dpi | | | | | |
|---|---|---|---|---|---|
| Sample Comparisons | Mean | Sample Size (n) | Sample Variance (s) | P(T<=t) first-tail | P(T<=t) second-tail |
| No Infection vs Infection only | n1 = 9.86 | 5 | 87.9 | 0.184 | 0.369 |
| | n2 = 15.2 | 5 | 68.5 | | |
| No Infection vs Infection ZA inject | n1 = 9.86 | 5 | 87.9 | 0.142 | 0.283 |
| | n2 = 16.4 | 5 | 75.2 | | |
| No Infection vs Infection AB inject | n1 = 9.86 | 5 | 87.9 | 0.174 | 0.349 |
| | n2 = 15.5 | 5 | 71.7 | | |
