## Supplementary material for "Functional genome and microbiome in blood of goats affected by the gastrointestinal pathogen *Haemonchus contortus*": Table 14. Statistical comparison of Firmicutes of 21 dpi

**Table 14. Statistical comparison of *Firmicutes* of 21 dpi.** Statistical comparison of *Firmicutes* (% Composition) after 21 dpi of host *Capra hircus* wethers.

| *Firmicutes* (% Composition) 21 dpi versus 21 dpi | | | | | |
|---|---|---|---|---|---|
| Sample Comparisons | Mean | Sample Size (n) | Sample Variance (s) | P(T<=t) first-tail | P(T<=t) second-tail |
| No Infection vs Infection only | n1 = 4.31 | 5 | 15.1 | 0.0618 | 0.1237 |
| | n2 = 9.87 | 5 | 35.4 | | |
| No Infection vs Infection ZA inject | n1 = 4.31 | 5 | 15.1 | **0.0368** | 0.0736 |
| | n2 = 11.1 | 5 | 36.3 | | |
| No Infection vs Infection AB inject | n1 = 4.31 | 5 | 15.1 | 0.076 | 0.153 |
| | n2 = 11.2 | 5 | 73.2 | | |
