## Supplementary material for "Functional genome and microbiome in blood of goats affected by the gastrointestinal pathogen *Haemonchus contortus*": Table 15. Statistical comparison of F/B ratios

**Table 15. Statistical comparison of *F*/*B* ratios.** Statistical comparison of *F*/*B* ratios % Composition after 21 dpi in host *Capra hircus* wethers.

<table>
<tr><th colspan="6">F/B ratio 21 dpi versus 21 dpi</th></tr>
<tr><th>Sample Comparisons</th><th>Mean</th><th>Sample Size (n)</th><th>Sample Variance (s)</th><th>P(T<=t) first-tail</th><th>P(T<=t) second-tail</th></tr>
<tr><td rowspan="2">No Infection vs Infection ZA inject</td><td>n1 = 0.606</td><td>5</td><td>0.044</td><td rowspan="2">0.015</td><td rowspan="2">0.030</td></tr>
<tr><td>n2 = 0.898</td><td>5</td><td>0.010</td></tr>
<tr><td rowspan="2">No Infection vs Infection AB inject</td><td>n1 = 0.606</td><td>5</td><td>0.044</td><td rowspan="2">0.035</td><td rowspan="2">0.070</td></tr>
<tr><td>n2 = 0.948</td><td>5</td><td>0.085</td></tr>
</table>
